# Intracellular acidification is a hallmark of thymineless death in *E. coli*

**DOI:** 10.1101/2022.04.04.487045

**Authors:** Alexandra Ketcham, Peter L. Freddolino, Saeed Tavazoie

## Abstract

Thymidine starvation causes rapid cell death. This enigmatic process known as thymineless death (TLD) is the underlying killing mechanism of diverse antimicrobial and antineoplastic drugs. Despite decades of investigation, we still lack a mechanistic understanding of the causal sequence of events that culminate in TLD. Here, we used a diverse set of unbiased approaches to systematically determine the genetic and regulatory underpinnings of TLD in *Escherichia coli*. In addition to discovering novel genes in previously implicated pathways, our studies revealed a critical and previously unknown role for intracellular acidification in TLD. We observed that a decrease in cytoplasmic pH is a robust early event in TLD across diverse genetic backgrounds. Furthermore, we show that acidification is a causal event in the death process, as chemical and genetic perturbations that increase intracellular pH substantially reduce killing. We also observe a decrease in intracellular pH in response to exposure to the antibiotic gentamicin, suggesting that intracellular acidification may be a common mechanistic step in the bactericidal effects of other antibiotics.

## Introduction

Deoxythymidine triphosphate (dTTP) is one of the four nucleoside triphosphates required for DNA replication. Its *de novo* biosynthesis requires the action of thymidylate synthetase, encoded by *thyA* in *E. coli*. When a *thyA* mutant is starved of exogenous thymidine, it rapidly dies [1, 2]. This killing phenomenon, TLD, was subsequently shown to occur in other bacteria, yeast, and human cells [3]. Thymidine starvation-induced killing is closely linked to the mode of action of several clinically important antibacterial, antineoplastic, and antimalarial drugs such as methotrexate [4], trimethoprim [5], and 5-fluorodeoxyuridine [6]. Resistance has been observed for all of these drugs [7–9]. Thus, a better understanding of the toxic phenomenon is likely to have broad clinical implications.

Although DNA synthesis is sharply inhibited during TLD, there is still an increase in DNA content in thymidine-starved *E. coli* cells [2, 10–12] and a correlation exists between the extent of lethality and the number of replication forks per cell [13]. TLD is reduced by mutations in or near the origin of replication (*oriC*) that interfere with the required transcription step for initiating replication [14, 15]. Inactivating the major DNA replication and repair proteins also modulates TLD kinetics, and the proteins involved fall into two categories: those with protective roles and those that enhance killing [3].

Early researchers in the field reported that stopping respiration (by oxygen removal) stops killing [16]. Several years later, it was discovered that *E. coli c*ells deficient in the respiration protein cytochrome oxidase, encoded by *cydA,* are protected from TLD in rich medium [17]. Recently it was found that the enhanced survival of respiration mutants is associated with lower accumulation of endogenous reactive oxygen species (ROS) levels, suggesting that it is the buildup of ROS and resultant damage to DNA that kill cells during thymidine starvation [18].

Although replication initiation, DNA damage, and ROS accumulation have all been observed in bacteria during thymidine starvation and implicated in the killing process, the precise sequence of events from thymidine starvation, DNA damage, and ROS accumulation to death remains unknown. Another central question is whether additional pathways also contribute to the killing process.

In this work, we undertook three complementary systems biology approaches to systematically probe the underlying mechanisms of TLD. We quantified the contribution of every non-essential gene in the genome to TLD, evolved strains that exhibit extreme TLD-resistance, and probed the transcriptional responses of TLD-sensitive and resistant strains. Genes involved in DNA replication and repair, electron transport chain and/or ROS accumulation, and pH homeostasis showed consistent effects across independent approaches and genetic backgrounds.

We observed that intracellular pH (pHi) strongly correlates with survival at both early and late time points, and that cytoplasmic acidification precedes ROS accumulation. Consistent with a causal role in death, experimental modulation of pH significantly affected survival. Finally, we provide evidence that intracellular acidification also occurs upon exposure to the antibiotic gentamicin, suggesting that it may be a common early step in the lethality of other bactericidal perturbations.

## Results

### Tn-seq Profiling Reveals a Role for pH Modulators in TLD

A saturated Tn5 transposon insertion library generated in a *thyA^-^* strain was selected in thymidine-free media and insertions were mapped and sequenced at different time points to look for changes in survival caused by gene disruptions (Fig 1A; S1A Fig). A survival score for each gene was defined as the log_2_ fold change (LFC) of fragments per million (FPM) at 3 hours (h) versus FPM at 0h (S1 Table).

**Fig 1.**
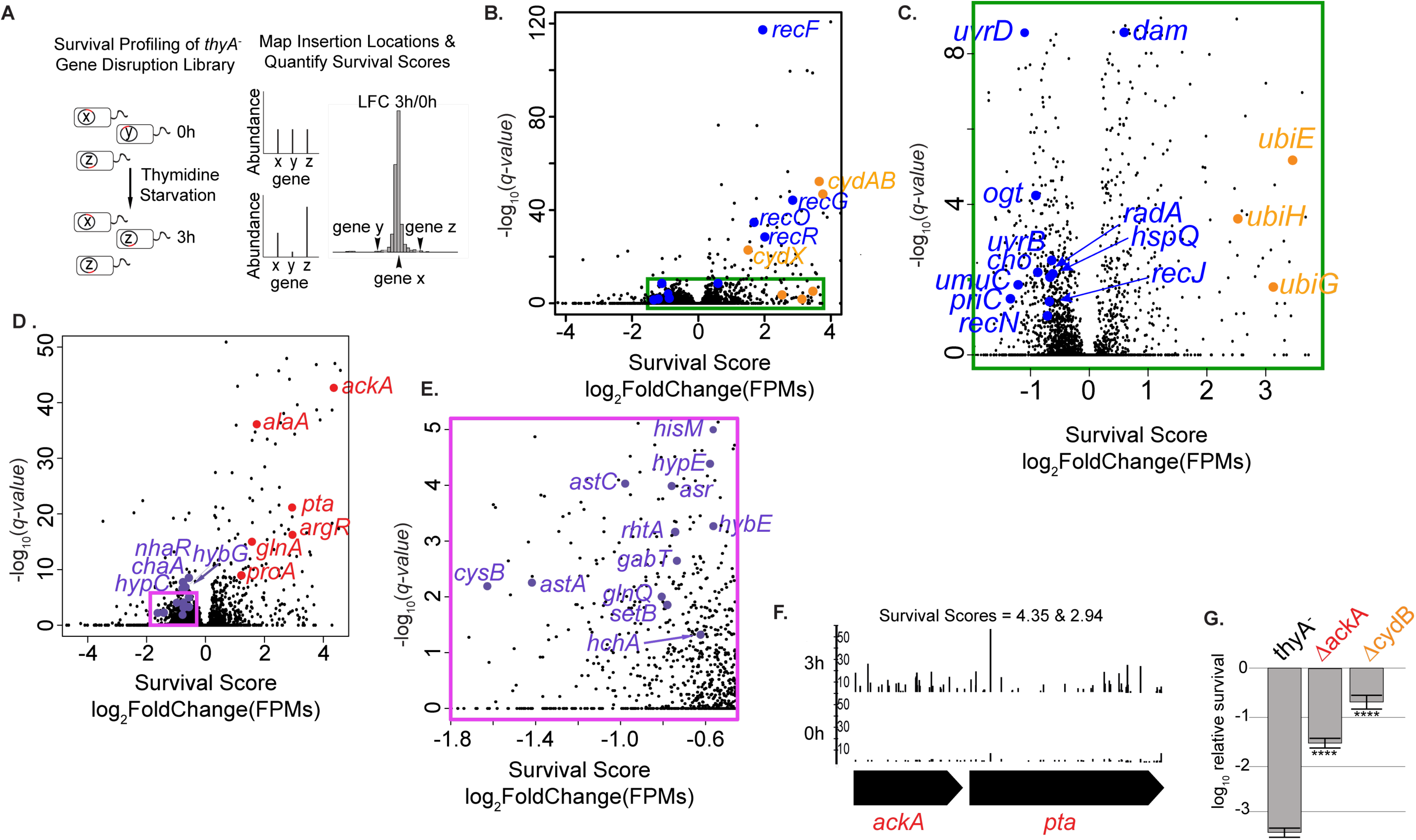
Systematic genetic survey of TLD by Tn-seq reveals a novel role for pH homeostasis. (A) A transposon insertion library was generated in the MG1655 *thyA^-^*strain and starved for thymidine. DNA adjacent to each insertion was amplified and sequenced at the beginning of the selection and at 3h. For each gene, the survival score is the log_2_ fold change of normalized reads at 3h/0h. (B-C) Volcano plot of survival score and significance showing genes in previously known pathways in color. Here and in the rest of this work, orange represents genes associated with respiration/ROS and blue represents genes involved in DNA replication/repair. Significance was calculated using an exact rate ratio test (using the rateratio.test R package) with a Benjamini & Yekutieli correction [19]. (C) is an inset for the area in the rectangle pictured in (B). (D-E) Volcano plot of survival score and significance showing in color genes from candidate lists that modulate cytoplasmic pH. Disruptions in genes involved in proton influx, or in lowering substrates needed for deacidification, have positive survival scores (shown in red). Conversely, disruptions in genes whose products are directly involved in proton consumption or producing substrates required for deacidification systems have negative survival scores (in light blue). (E) is an inset for the area in the rectangle pictured in (D). (F) Frequency of transposon insertions along the length of two newly identified TLD contributors at 0h and at 3h thymidine starvation. The enhancements at 3h are visualized using the Integrated Genome Browser [20]. The frequencies of each insertion site are shown in read counts per million. (G) Enhanced survival of validated genes. Candidates were validated by transferring Keio collection knockout alleles into MG1655 *thyA^-^*, and assessing their survival at 3h of thymidine starvation. All death assays, unless otherwise stated, were performed at 37°C. Relative survival was measured for at least three independent experiments, with error bars representing standard error of the mean. *p*-values for all death assays were calculated using a Welch t-test. * P<0.05, ** P<0.01, *** P < 0.001, **** P<0.0001. Here and in the rest of this work, red color represents genes in the acetate dissimilation pathway.

Tn-seq analysis revealed 212 genes in which insertions exacerbate killing and 175 genes in which insertions alleviate killing (S2 and S3 Tables; S1B-C Figs; Methods). As expected, disruptions in genes whose inactivation have previously been shown to sensitize cells to TLD, such as *uvrD* [21–23], have significant negative survival scores, and disruptions in genes whose inactivation have previously been shown to enhance survival, such as *recO* [22, 24, 25], have significant positive survival scores (Figs 1B-C, S1D Fig). Twenty additional genes with significant fitness effects fall in the previously known pathways of DNA replication and repair, and respiration (Figs 1B-C; Table 1). Over half of these genes have not been previously implicated in TLD to our knowledge.

**Table 1:** Genes from survival profiling candidate lists that fall into the previously known pathways of DNA replication/repair and respiration.

| Gene | Survival Score |
| --- | --- |
| *cho* | -1.21 |
| *cydABX* | 3.76, 3.65, 1.50 |
| *dam* | 0.59 |
| *hspQ* | -0.67 |
| *nth* | -0.71 |
| *ogt* | -0.90 |
| *priC* | -1.0 |
| *radA* | -0.62 |
| *recFGJNOR* | 1.95, 2.85, -0.64, -0.68, 1.68, 2.01 |
| *ubiEGH* | 3.46, 3.14, 2.53 |
| *umuC* | -1.34 |
| *uvrB* | -0.88 |
| *uvrD* | -1.1 |

A common theme among many of the genes identified by our Tn-seq survey, which do not belong to already known pathways, was involvement in pH homeostasis. We observed that disruptions in genes that produce or import H^+^ into the cytoplasm, or that lower levels of substrates needed for deacidification systems, enhance survival during thymidine starvation, and disruptions in genes that consume protons, or produce substrates needed for deacidification systems, exacerbate killing (Fig 1D-E and Table 2).

**Table 2:** Genes from survival profiling candidate lists that fall in the newly identified pathway of pH homeostasis.

| **Pathway** | **Gene** | **Survival Score** |
| --- | --- | --- |
| acetate dissimilation | *ackA* | 4.35 |
| glutamate metabolism | *alaA* | 1.74 |
| negative regulation of arginine biosynthesis | *argR* | 2.95 |
| acid shock protein | *asr* | -0.76 |
| arginine catabolic process to glutamate | *astA* | -1.42 |
| arginine catabolic process to glutamate | *astC* | -0.97 |
| sodium ion: proton antiporter | *chaA* | -0.76 |
| controls arginine-dependent acid resistance | *cysB* | -1.63 |
| glutamate formation | *gabT* | -0.73 |
| glutamate metabolism | *glnA* | 1.57 |
| L-glutamine transport | *glnQ* | -0.81 |
| response to acidic pH; mutant more sensitive to acid stress | *hchA* | -0.61 |
| lysine arginine uptake | *hisM* | -0.56 |
| hydrogenase 2-specific chaperone | *hybE* | -0.57 |
| hydrogenase 3 maturation protein | *hypC* | -0.78 |
| Na<sup>+</sup>/H<sup>+</sup> antiporter Regulator | *nhaR* | -0.57 |
| proline biosynthesis from glutamate | *proA* | 1.21 |
| acetate dissimilation | *pta* | 2.94 |
| proton antiporter; pH homeostasis | *rhtA* | -0.74 |
| proton antiporter | *setB* | -0.78 |

Insertions in genes *cydA* and *cydB* led to the highest survival scores in the list of candidates in previously known pathways (S1E Fig). The novel candidate contributor, *ackA,* encoding acetate kinase of the acetate production and excretion pathway, had the highest survival score of the candidate genes with a role in pHi modulation (Fig 1F). Acetate production and excretion play prominent roles in intracellular acidification since acetate, once excreted, can freely permeate back across the cell’s membrane into the cytoplasm where it dissociates delivering protons and lowering the pH [26, 27]. Inactivation of the *ackA* gene and the related gene in this pathway, *pta,* has been shown to cause constitutive extreme-acid resistance as well as less acidification in the medium [28, 29]. The effects of *cydB* and *ackA* on TLD were each validated by transferring the corresponding Keio knockout allele to the MG1655 *thyA^-^* strain and assessing their survival during thymidine starvation. Clean deletions in both of these genes showed significant enhancements in survival compared to the *thyA^-^* strain background (Fig 1G).

### Laboratory Evolution of Extreme TLD Resistance

A complementary and unbiased way to probe the *E. coli* genome for genes involved in TLD is through long-term laboratory evolution [30]. Clones of the *thyA^-^*strain in two genetic backgrounds (MG1655 and MDS42 [31]) were selected independently for increased survival in supplemented defined media in limiting amounts of thymidine (0.2μg/mL) and propagated for ∼50 days, with one transfer per day into fresh medium (Fig 2A). While each parental strain shows 2-3 orders of magnitude of death at 3h thymidine starvation, the best surviving isolate from each background shows very little death (Fig 2B). In fact, it takes the best surviving evolved isolates in each background 18-50h to suffer the same magnitude of loss as that suffered by the parental strain at 3h (Fig 2C).

**Fig 2.**
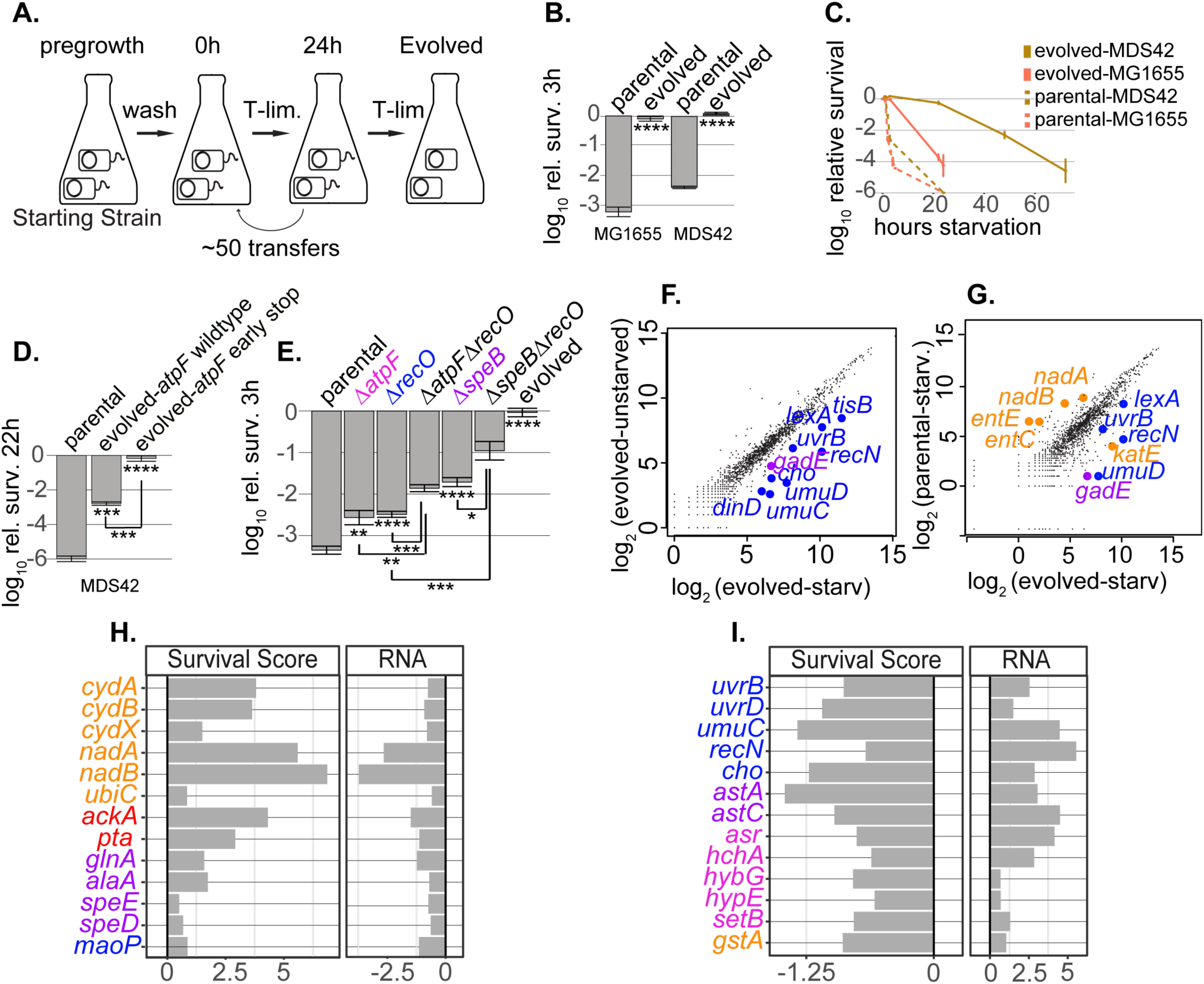
Genetic and transcriptional evidence for the role of pH homeostasis in laboratory evolution of extreme TLD-resistance. (A) Experimental setup of laboratory evolution. Clones of *thyA^-^*strains in two genetic backgrounds were selected independently in 0.2μg/mL thymidine. After roughly 50 transfers, isolates were characterized for enhanced survival. (B) Short-term death assay for evolved isolates and parents at 3h thymidine starvation. The stars in Fig 2 correspond to the *p*-values as specified in Fig 1 and are calculated using a Welch t-test.(C) Long-term death assay for evolved isolates and parents. The points are averages from independent experiments, with error bars representing standard error of the mean. (D-E) Survival of TLD-sensitive versus various TLD-resistant strains. The stars at the bottom of a bar represent the *p*-values calculated for that strain compared to the parental strain. The stars under brackets represent the *p*-values calculated between the strains bracketed. (D) *atpF* early stop contributes to extreme TLD resistance. Two sibling isolates from the same evolved population share all mutations except an *atpF* early-stop. The three strains shown are in the MDS42 background. (E) Clean deletion validations at 3h thymidine starvation. All knockouts are in the MG1655 *thyA^-^* background. Here and in the rest of this work, purple represents genes in putrescine / glutamate / arginine metabolism, and pink represents genes involved in proton translocation (or sequestration) systems. (F-G) Transcriptome profiling of TLD-sensitive and TLD-resistant strains. RNA was extracted and sequenced 30 minutes into thymidine starvation for the evolved strain in the MDS42 background and its parent strain. RNA was also extracted and sequenced in the unstarved condition. Select genes showing significant differential expression are shown in color. Rockhopper [32, 33] uses a negative binomial distribution to estimate the uncertainties in the read counts. The *p*-values are FDR-corrected using the Benjamini-Hochberg procedure [34]. (F) Gene expression of the evolved strain during starvation versus unstarved. (G) Gene expression of evolved versus parental strain during thymidine starvation. (H-I), Select genes with consistent effects across genetic backgrounds and experimental approaches. In each, the left-hand column shows survival scores from the survival profiling experiment in the MG1655 strain and the right- hand column shows LFC of mRNA expression (TPM) in the evolved versus parental in the MDS42 background during thymidine starvation. (H) Genes that exacerbate TLD. The left-hand-column shows positive survival scores and the right-hand column shows down-regulation of RNA expression levels of the evolved strain vs. parental during starvation. (I) Genes that alleviate TLD. The left-hand-column shows negative survival scores and the right-hand column shows up- regulation of RNA expression levels of the evolved strain versus parental during starvation. See S7 Table for the full list of genes showing consistent effects.

Whole-genome sequencing revealed that the number of mutations in the two evolved isolates ranged from 15 in the MDS42 evolved strain to 69 in the MG1655 evolved strain (Table 3 and S4 Table). Three of the mutations occurred in loci that were previously characterized in the TLD literature: *recO* [22, 24, 25] in the MG1655 evolved strain, and both *recJ* [23–25, 35], and *oriC* [15] in the MDS42 evolved strain.

**Table 3:**
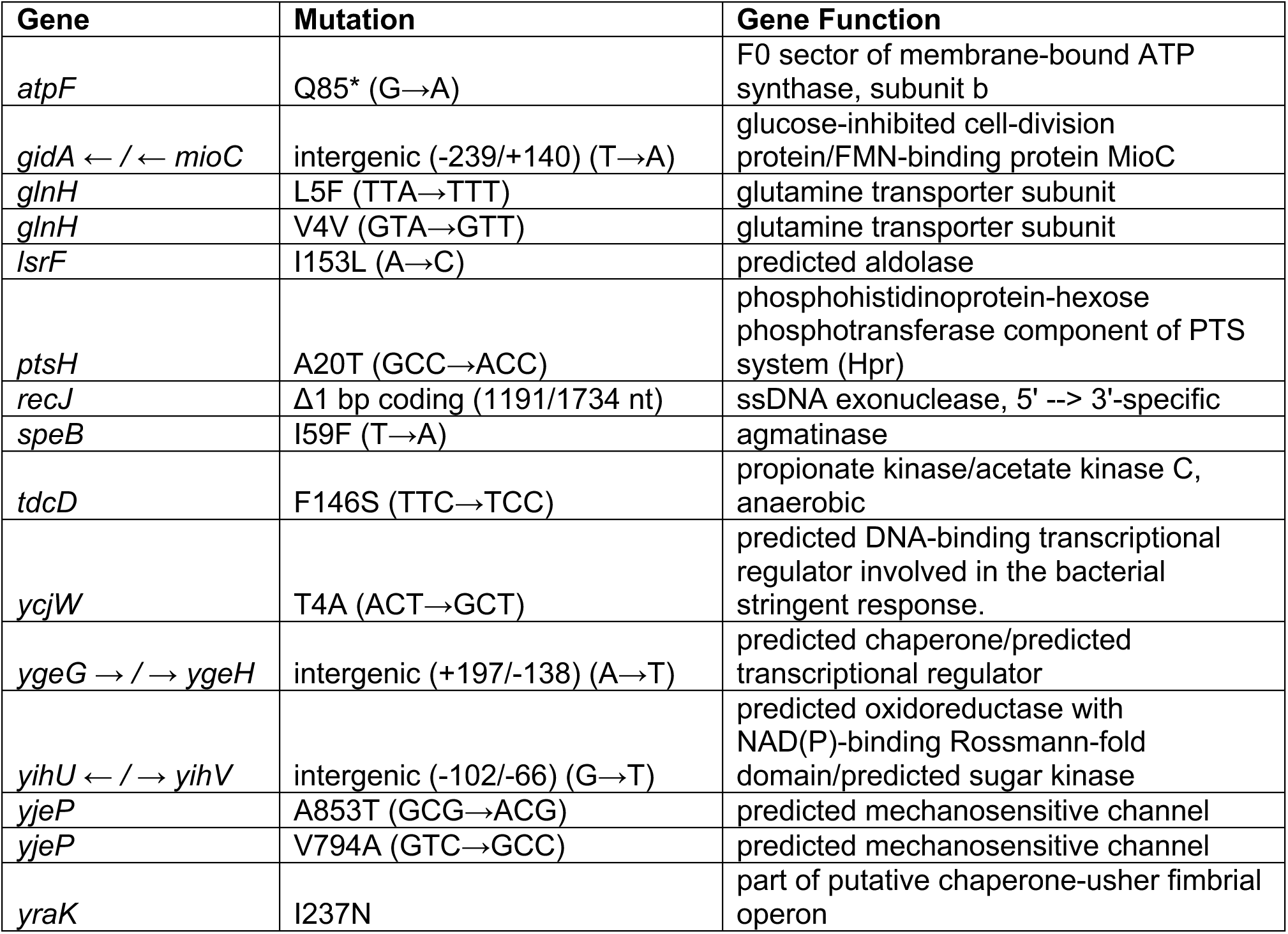
Mutations in the TLD-resistant evolved isolate in the MDS42 background.

The *oriC* and *recJ* mutations in the MDS42 evolved isolate are not causing all of this strain’s resistance. Another isolate derived from the same evolved population does not survive as well during thymidine starvation (Fig 2D). Whole-genome sequencing on both sibling isolates shows that they share all mutations except that the better surviving isolate has an *atpF* early stop codon. The *atpF* gene encodes subunit *b* of the proton-transporting (F0) portion of membrane bound ATP synthase. This protein complex catalyzes the synthesis of ATP using the energy of an electrochemical ion gradient by moving protons into the cell. It has not been previously implicated in TLD, but in principle is connected both to pH homeostasis and aerobic respiration.

To quantify the degree to which *atpF* plays a role in TLD resistance, a *ΔatpF* allele was transferred into the MG1655 thyA^-^ strain. Deletion of *atpF* significantly enhances the survival of the parental strain (Fig 2E) without slowing the growth rate in our defined media (Table 4, S2 Fig, and methods). Mutations in (or upstream of) the operon for the proton-transport-coupled ATP-synthase genes occur across both independently evolved isolates (Table 3 and S4 Table). A *recO* deletion was made in order to compare its survival with the novel mutants. Combining *ΔatpF* and *ΔrecO* significantly enhances the survival relative to either in isolation (Fig 2E). This increase in survival cannot be attributed to slower replication, as *ΔatpFΔrecO* has a faster growth rate than either mutation in isolation (Table 4 and S2 Fig).

**Table 4:**
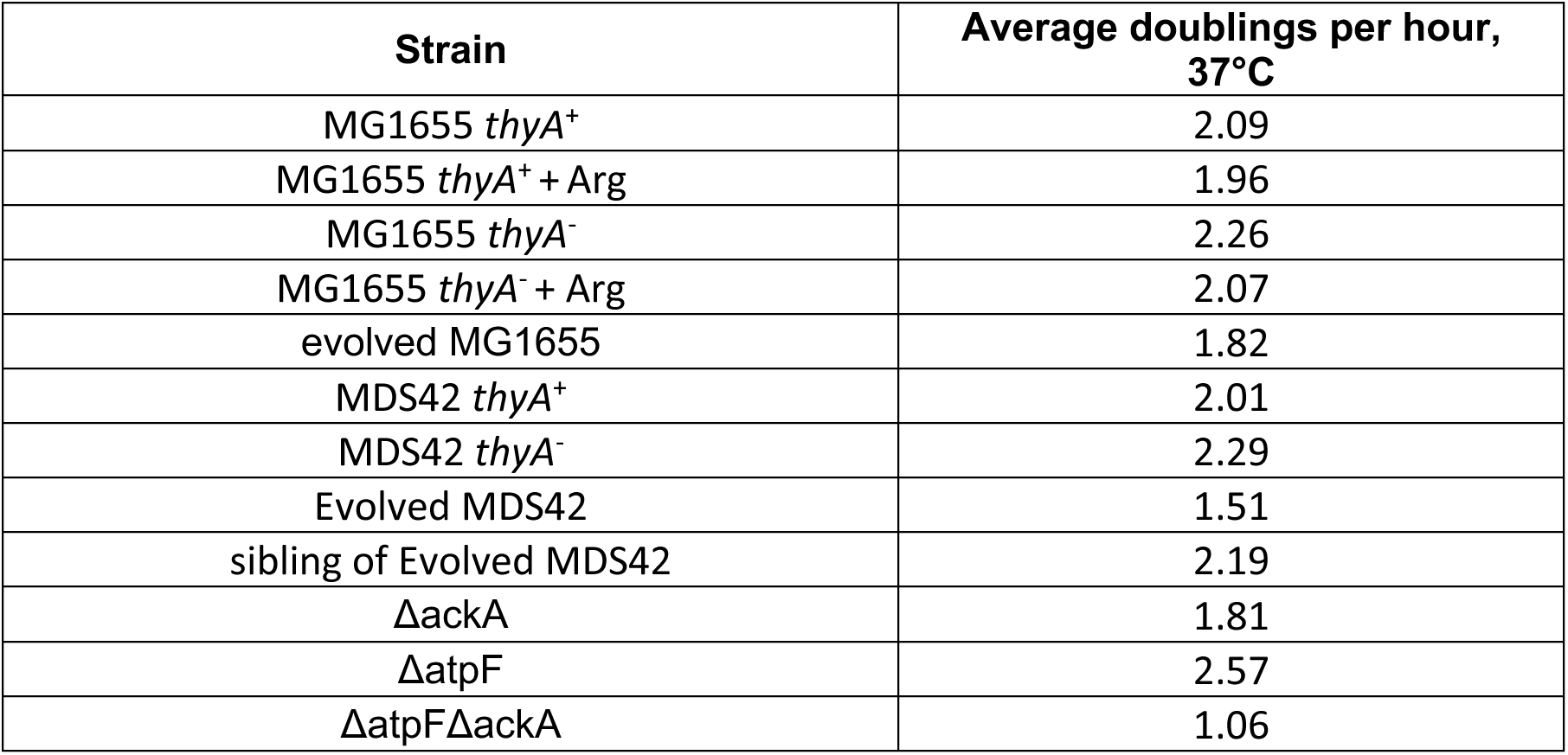
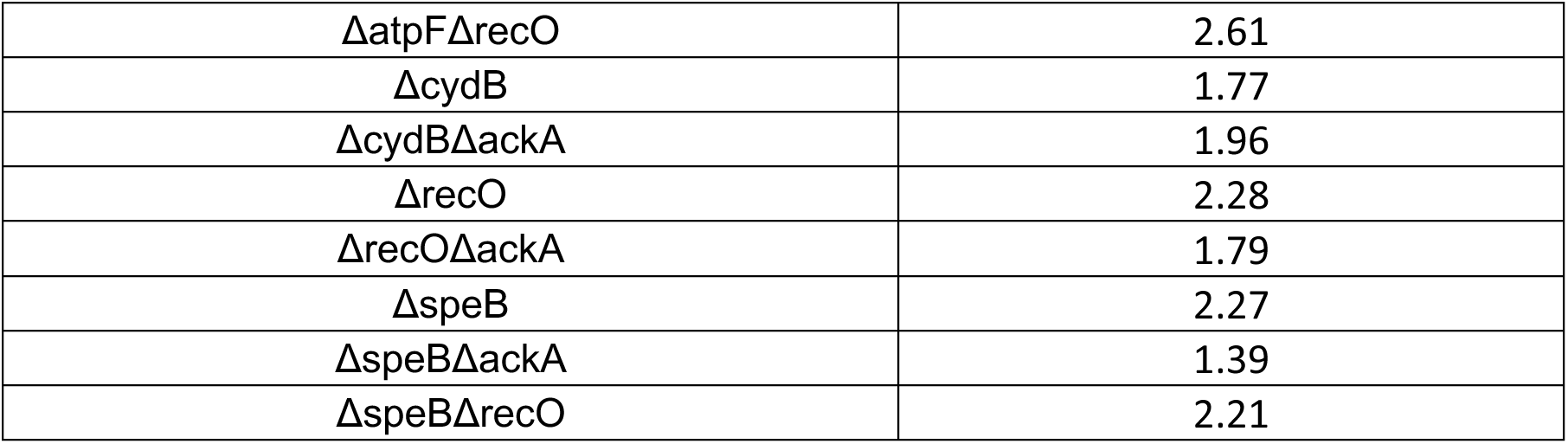
Growth rates of strains in rich defined media.

| Strain | Average doublings per hour, 37°C |
| --- | --- |
| MG1655 *thyA*<sup>+</sup> | 2.09 |
| MG1655 *thyA*<sup>+</sup> + Arg | 1.96 |
| MG1655 *thyA*<sup>-</sup> | 2.26 |
| MG1655 *thyA*<sup>-</sup> + Arg | 2.07 |
| evolved MG1655 | 1.82 |
| MDS42 *thyA*<sup>+</sup> | 2.01 |
| MDS42 *thyA*<sup>-</sup> | 2.29 |
| Evolved MDS42 | 1.51 |
| sibling of Evolved MDS42 | 2.19 |
| $\Delta ackA$ | 1.81 |
| $\Delta atpF$ | 2.57 |
| $\Delta atpF\Delta ackA$ | 1.06 |
| $\Delta atpF\Delta recO$ | 2.61 |
| $\Delta cydB$ | 1.77 |
| $\Delta cydB\Delta ackA$ | 1.96 |
| $\Delta recO$ | 2.28 |
| $\Delta recO\Delta ackA$ | 1.79 |
| $\Delta speB$ | 2.27 |
| $\Delta speB\Delta ackA$ | 1.39 |
| $\Delta speB\Delta recO$ | 2.21 |

In addition to recurring mutations in proton-transport-coupled ATP-synthase, both independently evolved isolates have mutations in putrescine biosynthesis genes, including a non- synonymous mutation in *speB*, whose transposon disruption significantly enhanced survival (S1F Fig; Table 3). *speB* encodes agmatinase, a putrescine biosynthesis enzyme. Putrescine has many diverse functions in the cell, including a role in pH homeostasis [36, 37]. Its metabolism is intricately linked to that of glutamate and arginine. Together these interlinked processes comprise three of the cell’s amino-acid-dependent decarboxylase acid resistance systems, which play critical roles in deacidification in *E. coli* [38]. When a *speB* deletion was transferred to the *thyA^-^* MG1655 strain, it showed significant enhanced survival compared to its parent (Fig 2E). Furthermore, a *ΔspeB* mutant combined with *ΔrecO* shows significantly enhanced survival compared to either strain in isolation (Fig 2E).

It is important to note that in principle, slowed bacterial growth might be expected to account for some of the effects on survival observed in both the evolved strains and targeted gene deletions considered here. However, apart from strains containing *ΔackA* alleles (which will be discussed further below), in general we did not observe a strong correlation between growth rate and TLD survival (S2 Fig) – for example, deletions of *recO, speB,* or *atpF* all strongly enhanced survival without affecting growth rate, and evolved strains show no to minor loss of growth rate while at the same time showing massively improved survival of TLD.

### The evolved TLD-resistant strain shows adapted transcriptional responses to DNA damage, ROS, and acid stress

In order to better understand how the evolved strain may have rewired its transcriptional output to enhance survival, RNA sequencing was performed on the evolved and parental strains in both thymidine-starved and unstarved conditions (S3A Fig and S5 Table). Strains in the MDS42 background were used, due to the lower number of accumulated (and potentially non-TLD related) mutations in the evolved strain. Whereas both the parental and evolved strain showed a transcriptional response to thymidine starvation, the evolved strain’s response was more pronounced (S3B-C Figs).

Several SOS-induced genes were significantly upregulated in the evolved strain in multiple comparisons, suggesting that these genes may have protective roles during TLD (Figs 2F-G and S4B-C Figs, in blue; S6 Table). Previous results have shown that, during TLD, the SOS response is induced [22, 25, 39, 40]. Our observations support a protective role for a subset of SOS-induced genes during the early stages of thymidine starvation.

In addition to the marked DNA-damage response, we saw evidence that the TLD-resistant strains have acquired improved capacity to cope with ROS. For example, *katE,* one of the two catalases in *E. coli* which serve as the primary scavengers for hydrogen peroxide [41, 42], is significantly upregulated in the evolved versus parental in both starved and unstarved conditions (Fig 2G; S4A-B Figs). The *de novo* NAD biosynthesis genes *nadA* and *nadB* are significantly downregulated in the evolved versus parental comparison during thymidine starvation (Fig 2G and S4B Fig, in orange). These genes have not, to our knowledge, been previously implicated in TLD survival. However, the product of *nadB* was reported to be a predominant source of endogenous hydrogen peroxide under aerobic conditions [43]. The product of *nadA* has an oxygen-sensitive iron-sulfur cluster that is required for its activity [44, 45]. Iron-sulfur clusters are known targets and sources of ROS due to the resulting displacement of iron from the cofactor and the intimate connection between labile iron and ROS generation [46, 47]. Iron-acquisition genes *entE* and *entC* are also significantly downregulated in the evolved versus parental comparison during starvation (Fig 2G and S4B Fig). Conversely, *entE* and *entF* are significantly upregulated in the parental starved versus unstarved (S4D Fig).

These data suggest that the TLD-resistant strain shows improved responses to both DNA damage and ROS accumulation during thymidine starvation. The significant upregulation of the gene *gadE* in the evolved strain, across multiple comparisons, suggests that TLD-resistant strains may also be responding to acid stress during thymidine starvation (Figs 2F-G; S4B-C Figs). *gadE* encodes the most important regulatory component of the glutamate-dependent acid response (GAD) system [48]. The product of *gadE* controls the transcription of *gadA* and *gadB*, genes involved in pH homeostasis [49].

### Consistent genetic and transcriptional evidence for the role of pH Homeostasis Genes in TLD

We hypothesized that genes whose products alleviate killing during thymidine starvation would not only have a negative survival score when disrupted, but also that their expression would be upregulated in the TLD-resistant strain. Conversely, genes whose products exacerbate killing during TLD were expected not only to show positive survival scores when disrupted, but that their expression would be down-regulated in the TLD-resistant strain. A search for genes that show these consistent effects in genetic and transcriptional responses yielded several lists of candidates (S7-S9 Tables). All three lists contain dozens of genes involved in DNA replication and repair, the electron transport chain and/or ROS accumulation, and pH homeostasis (Figs 2H-I; S5A-B Figs).

Two examples of genes in the pH homeostasis pathway that show consistent effects across experimental approaches are *glnA* (Fig 2H) and *setB* (Fig 2I). Disruptions in *glnA* which consumes glutamate, needed for the GAD system, helps survival during T-starvation (+1.57 LFC of FPMs), and expression of this gene is downregulated (−1.2 LFC of TPMs) in the resistant strain. Disruptions in *setB*, which encodes a proton antiport, involved in exporting protons out of the cell (just as the GAD system does), hurts survival (−0.78 LFC of FPMs) and expression of this gene is upregulated (+1.3 LFC of TPMs) in the resistant strain.

### Cells undergoing TLD show intracellular acidification before evidence of ROS accumulation

In order to test whether pH plays a role in TLD, we utilized two independent pH-sensitive dyes, BCECF-AM and pHrodo Green AM, to monitor pHi dynamics by flow cytometry. The absorbance of the fluorescein derivative 2’,7’-Bis-(2-carboxyethyl)-5-(and-6-) carboxyfluorescein 4 (BCECF) is sensitive to pH; its pKa makes it ideal for reporting pH_i_ at physiological conditions [50]. pHrodo is a newer rhodamine-based probe that is weakly fluorescent at neutral pH but interacts with hydrogen ions resulting in fluorescence (ThermoFisher Scientific). The addition of acetoxymethyl (−AM) groups to both probes makes them membrane permeable to facilitate loading into cells. Once inside, hydrolysis of the acetyl ester linkage by enzymatic cleavage regenerates the less permeable compound, which is retained within the intracellular space whereupon its fluorescence intensity acts as an indicator of pHi.

We also used Peroxy Orange 1, a cell permeable boronate compound specifically activated by intracellular H_2_O_2,_ to monitor ROS generation [51, 52]. Upon reaction with H_2_O_2_, a highly fluorescent product is released and can be assayed by fluorescence imaging.

Parental strains in both backgrounds showed evidence of ROS accumulation as well as intracellular acidification at 3h thymidine starvation (Figs 3A-F and S6A-C Figs; Methods). These effects are starvation-specific; they were not observed in the *thyA^+^* strain grown in thymidine-free media nor in the parental strain in thymidine-supplemented media (Figs 3A,C&E).

**Fig 3.**
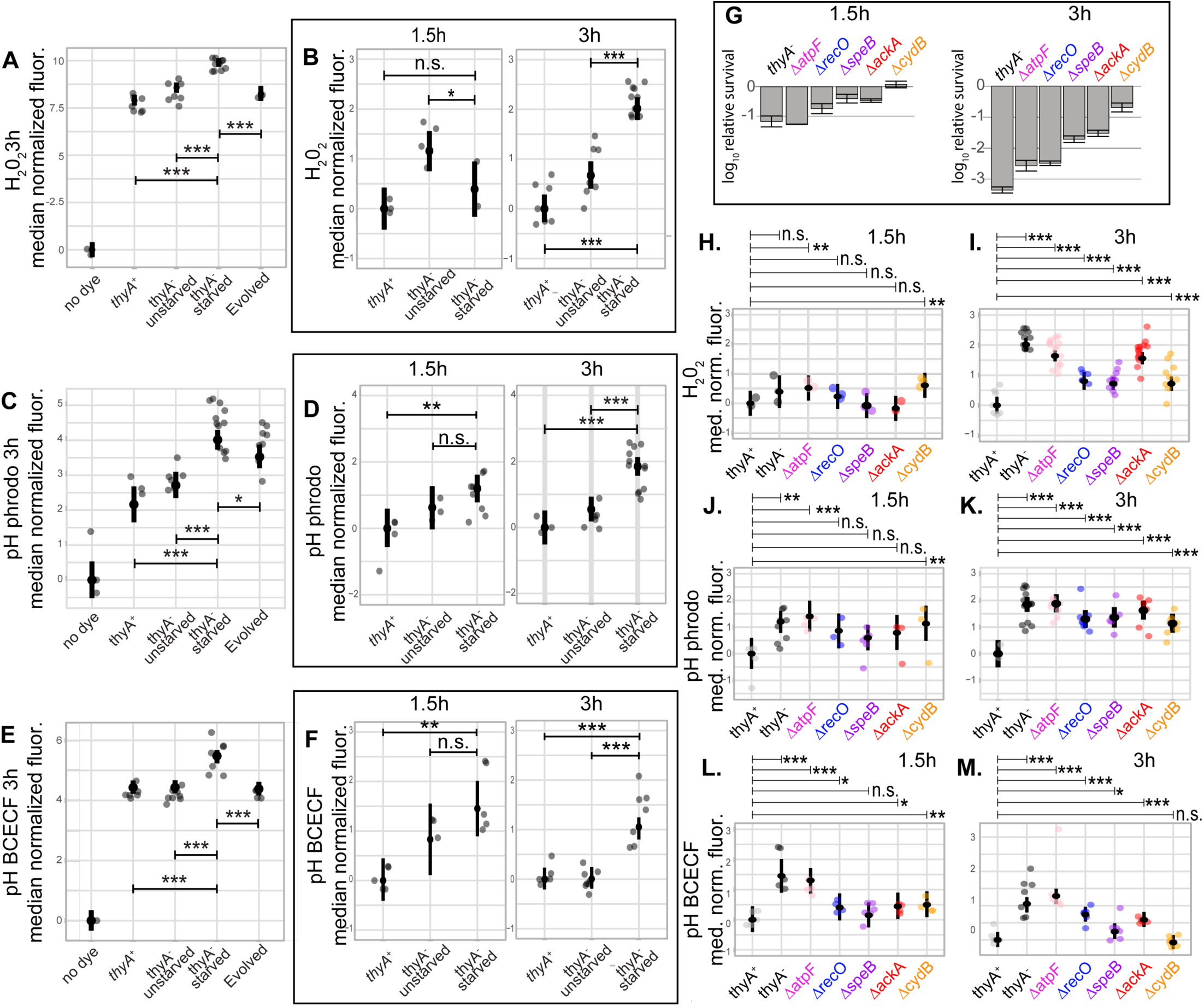
Cells undergoing TLD show intracellular acidification followed by ROS accumulation, with the degree of survival correlated with pH. Fluorescence data after accounting for cell size/shape (see Methods). Fluorescence was measured for each independent experiment using flow cytometry; in all flow cytometry figures unless otherwise noted, the large point for each condition shows the fitted effect for that biological condition, with error bars showing a 95% confidence interval. In addition, the median for each biological replicate is shown as a smaller translucent point (note that the shown biological replicates do not reflect corrections due to the random effect terms of the model, and thus show the actual biological variability of the experiment prior to model regularization). All values are offset by the fitted value for the first condition shown, which is thus centered on zero. Stars show significance tests based on the mixed effects model: * P<0.05, ** P<0.01, *** P < 0.001, **** P<0.0001. (A-F) Adjusted fluorescence for *thyA^+^*, parental, and evolved strains in the MG1655 background. All strains are in thymidine-free media except for those marked “unstarved”. (A-B) Adjusted fluorescence for strains dyed for H_2_O_2_ using Peroxy Orange. (A) Adjusted fluorescence measured at 3h. (B) Adjusted fluorescence measured at 1.5h and 3h. (C-D) Adjusted fluorescence measured for strains dyed for pHi using pHrodo Green. (C) Adjusted fluorescence measured at 3h. (D) Adjusted fluorescence measured at 1.5h and 3h. (E- F) Adjusted fluorescence measured for strains dyed for pHi using BCECF-AM. (E) Adjusted fluorescence measured at 3h. (F) Adjusted fluorescence measured at 1.5h and 3h. (G) Survival of the parental strain and knockouts in the MG1655 background at 1.5h and 3h thymidine starvation. (H-M) All strains shown are in thymidine-free media and in the MG1655 background. (H-I) Adjusted fluorescence for strains stained for H_2_O_2_ using Peroxy Orange. (H) Adjusted fluorescence at 1.5h thymidine starvation. (I) Adjusted fluorescence at 3h thymidine starvation. (J-K) Adjusted fluorescence for cells stained for pHi using pHrodo Green. (J) Adjusted fluorescence at 1.5h and (K) Adjusted fluorescence at 3h. (L-M) Adjusted fluorescence for cells stained for pHi using BCECF-AM. (L) Adjusted fluorescence at 1.5h. (M) Adjusted fluorescence at 3h.

While the parental strain showed acidification at 1.5h starvation for both pH dyes, it did not show evidence of ROS accumulation at this earlier time point, suggesting that acidification may precede ROS accumulation (Figs 3B,D&F). In order to ensure that the lack of observable ROS accumulation at the earlier time point was not the result of a delay in dye internalization or activation, the *thyA^-^* strain was visualized at 1.5h starvation after a 30-minute incubation with 1mM hydrogen peroxide. Since there is modest death at this early time point (Fig 3G at 1.5h), we assumed if ROS is involved it would be within the range of the lowest concentrations for mode one killing, which is due to DNA damage: 1 to 3 mM H_2_O_2_ [53]. We tested the lowest concentration within this range as in [54]. A significant elevation in ROS was observed for these samples (S7 Fig).

Unlike the TLD-sensitive strains, neither of the two evolved strains exhibited intracellular acidification nor showed signs of ROS accumulation at 3h thymidine starvation (Figs 3A,C&E and S6A-C Figs); likewise, the individual knockout strains that showed significant enhanced survival (Fig 3G) were also assessed for evidence of both ROS accumulation and intracellular acidification during thymidine starvation using flow cytometry, and showed general reductions particularly at 3h (Figs 3H-M).

### Manipulations that raise the pHi during thymidine starvation increase survival

Several of the genes that showed consistent effects across experimental approaches and backgrounds modulate intracellular levels of amino acids needed for the cell’s acid resistance decarboxylation systems. One of these amino acids, arginine, is a precursor of putrescine, and reductions in putrescine synthesis (such as those created by a *speB* knockout) have been observed to co-occur with increases in intracellular arginine concentration [55]. It is thus possible that one way in which the putrescine biosynthesis knockout enhances survival is by raising intracellular arginine levels. Shifts towards arginine accumulation would result in more robust intracellular deacidification, as the decarboxylation of arginine is one of the more powerful amino acid-dependent acid resistance systems in *E. coli* [56].

In order to test the direct involvement of pHi on TLD, L-arginine was added to growth media to increase the pHi and acid resistance [56, 57], and to see if there are resulting changes in survival. Indeed, adding 40mM L-arginine to *thyA^-^* strains, in both genetic backgrounds (MG1655 and MDS42), during thymidine starvation, substantially increased survival and this was accompanied by significant increases in pHi (Figs 4A-C). Adding 40mM L-arginine to *thyA^-^* strains in high thymidine media did not result in major growth defects that could account for these dramatic increases in survival (Table 4 and S2 Fig).

**Fig 4.**
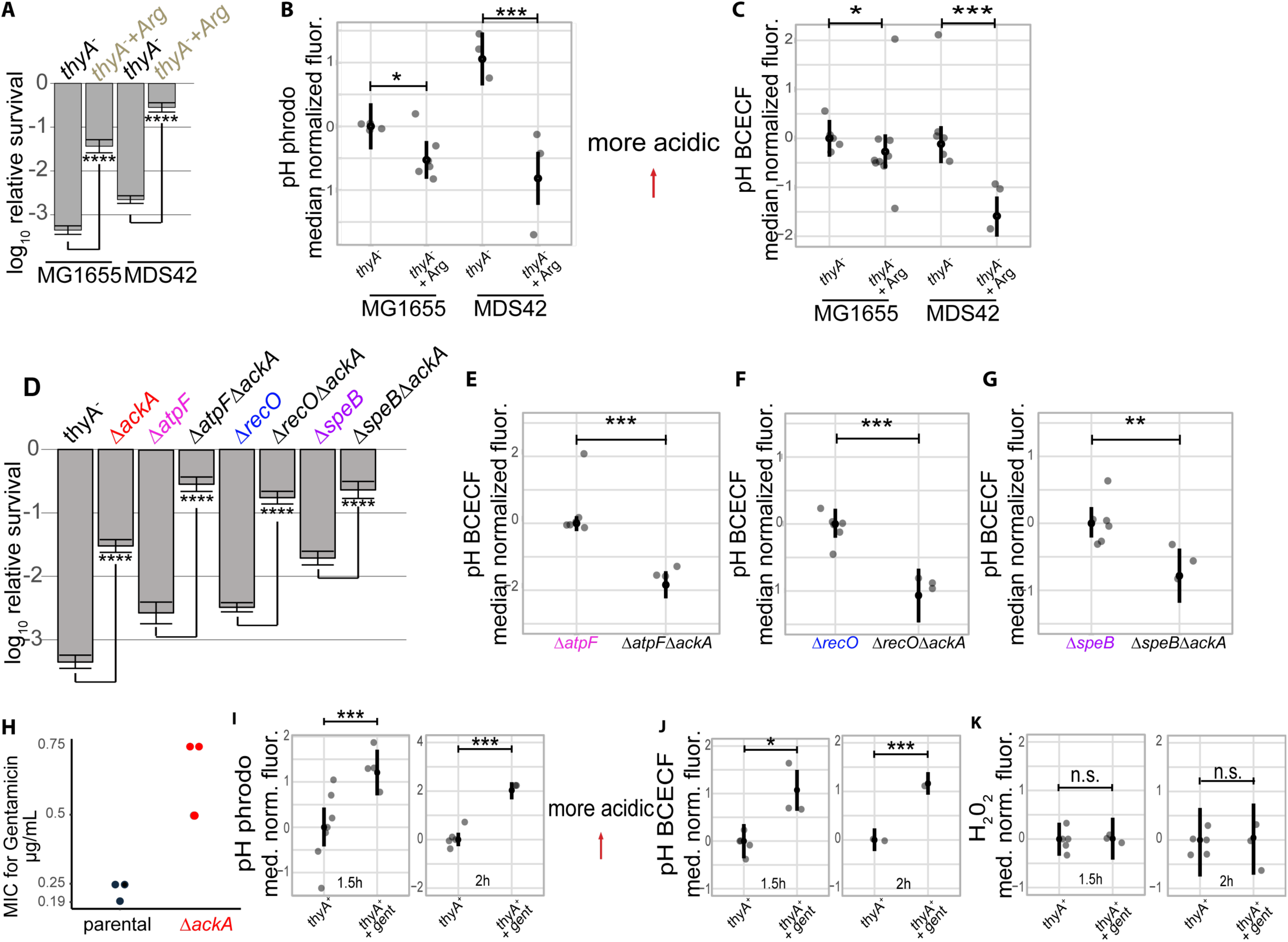
Experimental manipulation of pHi modulates survival. (A-C) Impact of arginine supplementation on survival and pHi. (A) L-arginine significantly increases the survival of *thyA^-^* strains in two backgrounds at 3h thymidine starvation. (B-C) Fluorescence was measured for each independent experiment for strains stained for pHi at 3h thymidine starvation and measured using flow cytometry using pHrodo Green (B) and BCECF-AM (C). (D-G) Impact of *ackA* deletion on survival and pHi. (D) Deletion of *ackA* from various resistant mutants results in significant increases in survival at 3h thymidine starvation. (E-G)) Adjusted fluorescence of strains stained for pHi at 3h thymidine starvation and measured using flow cytometry using BCECF-AM. (H) Minimum inhibitory concentration (MIC) of the *thyA^-^*and the Δ*ackA* strain for the antibiotic gentamicin. The concentrations are in μg/mL. (I-J) Adjusted fluorescence of wild type (*thyA^+^*) cells stained for pHi at 1.5h with and without 1μg/mL gentamicin treatment and measured using flow cytometry using pHrodo Green (I) or BCECF-AM (J). (K) Adjusted fluorescence of wild type (*thyA^+^*) cells stained for ROS accumulation at 1.5h with and without 1μg/mL gentamicin treatment and measured using flow cytometry using Peroxy Orange. The *p*-values in (A) and (D) were calculated as for the death assays in the rest of this work, using a Welch t-test. See Fig. 3 caption for definitions of plotted values and significance tests for flow cytometry data.

Genetic deletions of the acetate metabolism genes *ackA* and *pta* have been shown to enhance *E. coli’s* acid resistance [28, 29]. Removal of the *ackA* gene and the subsequent decrease in acetate production and excretion result in less acetate permeating back across the cell membrane and delivering protons into the cytoplasm [58]. Deletion of *ackA* from the *ΔatpF, ΔrecO*, and *ΔspeB* strains results in significant alleviation of TLD (Fig 4D). The increased survival of each was accompanied by significant elevations in pHi during thymidine starvation (Figs 4E-G). Deletion of *ackA* from the *ΔcydB* mutant resulted in no significant increase in survival (S8 Fig; see Discussion).

Part of the effects that we observe from deletion of *ackA* likely arise due to the decreased growth rate of *ΔackA* cells in our experimental conditions, but changes in growth rate do not fully explain the protection from TLD. We found that the *thyA* mutant grown at 30° has a lower growth rate than either *ΔackA* or *ΔrecO ΔackA* at 37° (S2 Fig), but that its growth at 30° had a much less substantial effect on survival than the additional mutations (S9 Fig). Also note that since the three double mutants with similar survival levels at 3h starvation have very different growth rates, changes in replication rates cannot account for all the increased survival associated with *ΔackA* cells (Table 4 and S2 Fig).

### Intracellular acidification occurs following exposure to gentamicin

To test whether the genetic manipulation of pHi by deletion of *ackA* increases the resistance to other bactericidal perturbations, we measured the minimum inhibitory concentration (MIC) of the *ΔackA* strain for various antibiotics. The MIC for gentamicin applied to the *ΔackA* strain was three-fold higher than for the corresponding parental strain (Fig 4H). In order to test whether pH changes also accompany exposure to gentamicin, we stained *thyA^+^*cells with pH-sensitive dyes, treated with a drug concentration that generated under 1 log-fold death at 1.5h, and looked for pH changes by flow cytometry. Both independent pH dyes showed significant acidification shifts 1.5h and 2h following drug exposure compared to the no-drug control (Figs 4I-J). We also stained these cells for ROS and observed no changes in ROS at 1.5h or 2h between the gentamicin treated cells and no-drug cells controls (Fig 4K). At higher concentrations of gentamicin, we saw both acidification and ROS changes (Fig S10). These data suggest that the pH drop that is a key contributor to the killing process during thymidine starvation independently of ROS also occurs in other bactericidal perturbations, and may play a broader mechanistic role in cell death.

## Discussion

We used a set of complementary systems biology approaches to probe the underlying mechanisms of TLD in *E. coli*. We quantified the contribution of every non-essential gene in the genome to TLD via Tn-seq, evolved strains that exhibit unprecedented long-term TLD-resistance, and probed the transcriptional responses of TLD-sensitive and resistant strains. Dozens of novel candidate genes were identified across the various approaches. Among the recurring biological processes that modulated survival was intracellular pH homeostasis. Multiple independent lines of evidence support a critical role for cytoplasmic acidification in TLD. All three novel validated beneficial loss-of-function mutations arising from our study, *ΔspeB, ΔackA,* and *ΔatpF*, are in pathways that influence cytoplasmic pH. Mutations in the putrescine biosynthesis pathway, and in genes in or upstream of the operon encoding ATP synthase, recurred in the two best-surviving independently evolved isolates in different genetic backgrounds (MG1655 and MDS42). TLD sensitive strains showed intracellular acidification during thymidine starvation. Survival of all of the tested strains strongly associates with intracellular acidification throughout the death process. Finally, manipulations that increased the pHi resulted in corresponding increases in survival during thymidine starvation—in some instances without significant changes in ROS accumulation.

Previous researchers have proposed, and our data supports, a model in which replication initiation during thymidine starvation causes DNA damage. A subset of SOS-induced genes are significantly upregulated in the TLD-resistant strain and show a survival cost when disrupted in the *thyA^-^* strain, confirming that some DNA repair genes have protective roles during thymidine starvation. Other DNA damage repair proteins, such as *recO,* aggravate the damage. The enhanced survival of *ΔrecO* cells is associated with both lower ROS levels and less intracellular acidification. Although *ΔrecO’s* involvement in TLD is not new, our finding that the knockout lowers both ROS levels and intracellular acidification suggests a potential link between DNA damage and ensuing acidification and ROS generation.

We propose that thymidine starvation results in three toxic outcomes in the cell: DNA damage, intracellular acidification, and ROS accumulation. We observe acidification before evidence of ROS accumulation. It is possible that ROS and acidification are two independent responses and acidification simply occurs earlier. It is also possible that acidification triggers ROS accumulation. The fact that targeted gene deletions that we identified in pHi-altering pathways also alleviate ROS accumulation during thymidine starvation (and especially that in some cases beneficial deletions affect TLD without bringing down ROS levels; Fig. 3H-M) argues for the latter interpretation.

Several groups have previously suggested that acid and oxidative stress can synergize. Many acid stress genes overlap with those of oxidative stress, and researchers have proposed that low pH amplifies the toxicity of radicals [59–61]. Glutamate and arginine have also been reported to play important roles in the protection against oxidative stress under acidic conditions in *E. coli* [62], partly through enhanced nitric oxide production, and thus contributions orthogonal to effects on pH cannot be ruled out. Oxidation reactions are affected by pH, so an increase in pH achieved via these acid resistance systems could modulate the outcome of oxidative reactions and processes [63, 64].

Both intracellular acidification and ROS can damage DNA. Deletion of *ackA* has been reported to have protective effects on DNA integrity [65]. Previous groups have shown that *ackA* and *pta* are associated with DNA replication and/or repair through as yet unknown mechanisms [65, 66]. It is possible that *ΔackA’s* disruption in acetate production somehow affects DNA replication and/or repair through a reduction in acidification. It is also possible that ROS accumulation and acidification contribute to TLD in ways separate from their causation of DNA damage.

The lack of increased survival in *ΔcydBΔackA,* relative to either mutant in isolation, is potentially informative. It has been reported that *ΔcydB* upregulates several acid resistance genes and exhibits acid resistance [67], and may explain the lack of acidification that we observed during thymidine starvation in this mutant. This group also reported that in *ΔcydB c*ells*, poxB* is upregulated [67]. *poxB* encodes pyruvate oxidase, the key enzyme in an alternate, low-efficiency pathway to synthesize acetate [68]. There have been two additional reports of increased activity of the PoxB pathway in a Δ*cydB*Δ*cyoB*Δ*nuoB* triple mutant [69, 70]. If *ackA* expression is already low in the *ΔcydB* strain in favor of *poxB* expression, removing *ackA* gene from the *ΔcydB* strain may result in minimal additional benefits. A potential role for the PoxB enzyme in alleviation of acidification and/or ROS accumulation during TLD is an interesting avenue for future research due to its ability to transfer electrons directly to the terminal oxidases and potentially uncouple respiration from ATP synthesis (and from the coincident proton influx) [71–73].

Several questions remain, particularly about the nature of the path that leads from thymidine starvation to acidification and (later) ROS accumulation. A recent study reported that trimethoprim treatment induces an acid stress response in *E. coli*; the researchers proposed that adenine depletion may cause a drop in pH [74]. Another group more recently linked antibiotic- induced adenine limitation to oxidative stress [75]. Thymidine starvation, DNA damage, and attempts to repair the damage, all cause nucleotide pool disruptions [76–79]. It is possible that dysregulation in adenine levels is the source of both the pH drop and the resulting increase in ROS levels during thymidine starvation. ATP levels may play an additional role in TLD independent of ROS and pH effects, and may help explain the enhanced survival of the *ΔatpF* strain.

Another outstanding question is whether TLD is an active or passive process. It is interesting to note that the transcription of siderophores involved in iron acquisition and ROS generation are significantly upregulated in the *thyA^-^* strain in the comparison of starved versus unstarved conditions and downregulated in the evolved starved versus unstarved conditions. This suggests possible triggering of an active ROS death pathway. Alternatively these processes may be activated in response to the pH drop in the TLD-sensitive strain. Previous work has shown that lowering of pH increases the availability of free iron [80].

A more parsimonious model is that TLD is a passive process and that ROS, acidification, and DNA damage constitute a perfect storm. Under normal conditions, drops in pH or spikes in ROS levels would cause only minor damage to DNA that the cell can recover from. However, when this damage is compounded by previous DNA damage caused by replication initiation and repair during thymidine starvation, the results are lethal. Regardless of whether TLD is a passive or active process, we conclude that intracellular acidification plays an instrumental role in the sequence of events from thymidine starvation to cell death. Our findings that genetic manipulations that increase pH also increase gentamicin resistance, and that exposure to gentamicin results in significant pH drops provide intriguing evidence that intracellular acidification may be a common causal factor in bacterial cell death caused by other bactericidal perturbations.

## Methods

### Strains and Culture

Thymidine auxotrophs were generated in two genomic backgrounds, MG1655, the prototype K-12 strain, and the “Multiple Deletion Series 42” (MDS42), a reduced-genome derivative of MG1655 [31] (See Table 5 for full list of strains used). The latter strain has 704 nonessential genes as well as insertion sequence elements and cryptic prophages deleted, amounting to a 14% reduction in the genome.

**Table 5:** Strains used in this work.

| Strain | Genotype | Source |
| --- | --- | --- |
| *thyA*<sup>+</sup> | MG1655 | MG1655 ATCC 700926 [81] |
| MG1655 *thyA*<sup>-</sup> | MG1655 *thyA*<sup>-</sup> | this work |
| MDS42 *thyA*<sup>+</sup> | MDS42 | [82] via Alison Hottes |
| MDS42 *thyA*<sup>-</sup> | MDS42 *thyA*<sup>-</sup> | this work |
| Δ*ackA* | MG1655 *thyA*<sup>-</sup> Δ*ackA* | this work |
| Δ*cydB* | MG1655 *thyA*<sup>-</sup> Δ*cydB* | this work |
| Evolved MG1655 | *thyA*<sup>-</sup> see S4 Table for list of mutations | this work |
| Evolved MDS42 | *thyA*<sup>-</sup> see Table 3 for list of mutations | this work |
| Sibling isolate of Evolved MDS42 | MDS42 *thyA*<sup>-</sup> all mutations in S6 Table except *atpF* is wild type | this work |
| Δ*atpF* | MG1655 *thyA*<sup>-</sup> Δ*atpF* | this work |
| Δ*recO* | MG1655 *thyA*<sup>-</sup> Δ*recO* | this work |
| Δ*atpF*Δ*recO* | MG1655 *thyA*<sup>-</sup> Δ*atpF* Δ*recO* | this work |
| Δ*speB* | MG1655 *thyA*<sup>-</sup> Δ*speB* | this work |
| Δ*speB*Δ*recO* | MG1655 *thyA*<sup>-</sup> Δ*speB* Δ*recO* | this work |
| Δ*atpF*Δ*ackA* | MG1655 *thyA*<sup>-</sup> Δ*atpF* Δ*ackA* | this work |
| Δ*recO*Δ*ackA* | MG1655 *thyA*<sup>-</sup> Δ*recO* Δ*ackA* | this work |
| Δ*speB*Δ*ackA* | MG1655 *thyA*<sup>-</sup> Δ*speB* Δ*ackA* | this work |
| Δ*cydB*Δ*ackA* | MG1655 *thyA*<sup>-</sup> Δ*cydB* Δ*ackA* | this work |

The *thyA* gene has a transcription terminator within the coding sequence that acts as the stop signal for the upstream essential gene *umpA* [83–85]. In order to remove t*hyA* function and preserve viability, we made an internal deletion without disrupting the overlapping transcript [84] via one-step inactivation using PCR products [86]. The *thyA* mutation in both starting strains is identical and has been verified with PCR, sequencing, and by failure to grow in thymidine-free media and agar plates. Both starting strains exhibit rapid and steep death in thymidine-free media.

Single gene deletions were obtained from the Keio collection [87] and transferred to the *thyA^-^* strain in the MG1655 background by P1 transduction [88] followed by selection on LB/ kanamycin/thymidine plates. Kanamycin-resistant clones were tested for the clean deletion by PCR, and then cured of the resistance cassette by transformation with the plasmid pcp20 [89] prior to characterization.

All strains in the *thyA^-^* background, unless they were being starved, were supplemented with 50μg/ml of thymidine (Sigma Aldrich, T1895) in liquid media (called “high thymidine” in this work) and all plates contained 15g/L agar and 50μg/ml of thymidine, unless otherwise stated.

For all experiments in liquid, strains were grown in defined supplemented M9 media either at 30°C or 37°C. In all experiments, unless otherwise stated, the liquid media was the same: 1x M9 salts (BD 248510) supplemented with a synthetic complete amino acid supplement (USBiological, D9515) that also supplies uracil and adenine; cytosine and guanine (each at 133μM); a vitamin solution (each in the 10μM-20μM range) of thiamine, folic, riboflavin and alpha lipoic acid; micronutrients: boric acid, cobalt chloride, copper sulfate, manganese chloride, zinc sulfate, ammonium molybdate, ferric citrate, MgS04, CaCl2 (each in the 10μM-10mM range, following the same formulation as in MOPS media [90]); and 0.2% glucose. Growth rates in high thymidine in our rich defined media were higher than in LB for every strain we tested (Table 6).

**Table 6:**
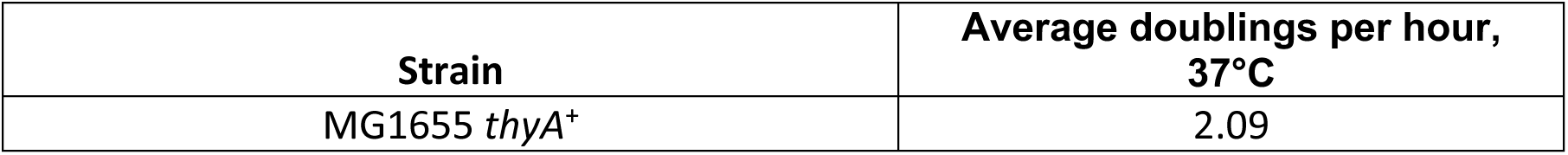
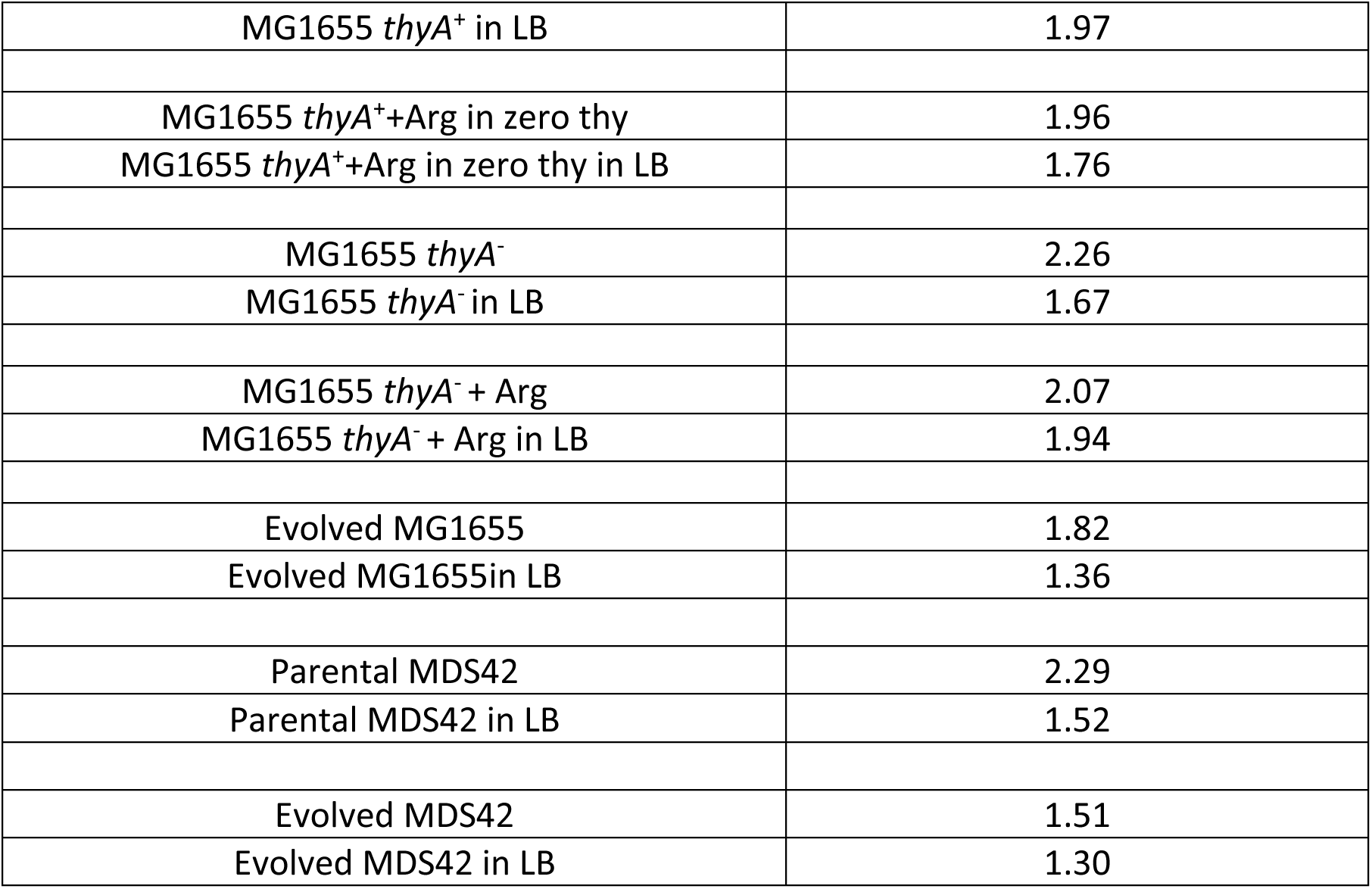
Growth rates of strains in rich defined media vs. LB.

| Strain | Average doublings per hour,<br>37°C |
| --- | --- |
| MG1655 *thyA*<sup>+</sup> | 2.09 |
| MG1655 *thyA*<sup>+</sup> in LB | 1.97 |
| MG1655 *thyA*<sup>+</sup>+Arg in zero thy | 1.96 |
| MG1655 *thyA*<sup>+</sup>+Arg in zero thy in LB | 1.76 |
| MG1655 *thyA*<sup>-</sup> | 2.26 |
| MG1655 *thyA*<sup>-</sup> in LB | 1.67 |
| MG1655 *thyA*<sup>-</sup> + Arg | 2.07 |
| MG1655 *thyA*<sup>-</sup> + Arg in LB | 1.94 |
| Evolved MG1655 | 1.82 |
| Evolved MG1655 in LB | 1.36 |
| Parental MDS42 | 2.29 |
| Parental MDS42 in LB | 1.52 |
| Evolved MDS42 | 1.51 |
| Evolved MDS42 in LB | 1.30 |

### Thymineless Death Assays

Overnight cultures of bacteria grown from single colonies were diluted 1:200 into 2mL high thymidine media and grown for 2h with shaking at 37°C. They were washed twice and placed in thymidine-free media, plated at 0h on LB/thymidine plates and again after the stated time of thymidine starvation with shaking at 37°C. The number of independent experiments for 3h Death assays: *thyA^-^* in MG1655: n=18; *ΔackA*: n=15; *ΔcydB*: n=9; evolved MG1655: n=7; *thyA^-^*in MDS42: n=9; evolved in MDS42: n=8; *ΔatpF*: n=8; *ΔrecO*: n=7; *ΔatpFΔrecO*: n=7; *ΔspeB*: n=11; *ΔspeBΔrecO*: n=7; *ΔatpFΔackA*: n=4; *ΔrecOΔackA*: n=6; *ΔspeBΔackA*: n=6. The number of independent experiments for the long-term death assay (Fig 2C): *thyA^-^* MG1655: (3h) n=18, (4h) n=3, (24h) n=1; evolved in MG1655: (3h) n=7, (22h) n=2, (24h) n=3; *thyA^-^* MDS42: (3h) n=9, (24h) n=3; evolved MDS4: (3h) n=8, (22h) n=3, (24h) n=3, (48h) n=3, (72h) n=2. The number of independent experiments for the 22h death assay (Fig 2D): *thyA^-^* in MDS42: n=3; *atpF* wild type (sibling isolate) strain: n=3; *atpF* early stop (evolved in MDS42): n=3.

#### Death assays with L-arginine supplementation

Overnight cultures were prepared as above. After 2 washes, cells were placed in thymidine-free media along with 40mM L-arginine, and plated for colony counting on LB/thymidine plates at 0h and again after 3h of thymidine starvation (with arginine) shaking at 37°C. For the arginine supplemented death assays: *thyA^-^* in MG1655: n=6; *thyA^-^* in MG1655 + Arg: n=4; *thyA^-^* in MDS42: n=3; *thyA^-^* in MDS42 + Arg: n=3; *ΔackA*: n=15; *ΔackA* + Arg: n=5.

#### Death assays with antibiotic treatment

After the pregrowth and washes as outlined above, thyA^+^ cells were treated with 1μg/mL gentamicin and plated for colony counting on LB/thymidine plates at 0h and again after 1.5h of shaking in defined media at 37°C. The number of independent experiments for this death assay at 1.5h: thyA^+^: n=4; thyA^+^+ gent1: n=5.

### Characterization of Growth Rates

Overnight cultures of bacteria grown from single colonies were diluted 1:200 in 150μl of fresh media and grown in a Biotek Synergy MX plate reader shaking continuously for up to 24h at 37°C. The absorbance at 600nm (OD 600) was measured every 10 minutes. Doubling times were calculated and provided in Table 4.

### Survival profiling

#### Transposon insertion library generation

Using Gibson assembly, we redesigned the EZ-Tn5 Transposon Vector (Epicentre) so that the transposon insertion has, in addition to the kanamycin cassette, an Illumina sequencing adapter. MG1655 *thyA^-^* cells were prepared for transformation and electroporated with 2μL of the generated transposomes. After 1h, serial dilutions were made of a small aliquot onto LB/kanamycin/thymidine plates in order to gauge the number of successful transformants. The remainder of the cell mixture was added to 250mL of LB/kanamycin/thymidine and grown to amplify the mutants. After 8h, the library was pelleted and resuspended in a small volume of 15% glycerol (diluted in LB), subsequently aliquoted, snap-frozen, and stored at −80°C. Our library had 174,101 insertions in the genome, or ∼1 insertion every 27 base pairs.

#### Selection of transposon library

The library was thawed and pre-grown for 2h in high thymidine, washed twice in 1x phosphate buffered saline (PBS), and placed in thymidine-free media. Once the library was placed in thymidine-free media, a “Start” sample was collected for DNA extraction. Another sample, “0h”, was taken at the same time but placed in high thymidine and allowed to undergo 2-3 doublings before DNA extraction. After 3h of thymidine starvation, a final sample, “3h” was placed in high thymidine for an outgrowth of 2-3 doublings before DNA extraction.

To summarize, the “Start” sample was taken at the very beginning of the thymidine starvation selection and was never placed in high thymidine for an outgrowth. Conversely, both “0h” and “3h” samples were thymidine-starved for the designated amount of time (0h and 3h) and an aliquot was placed in high thymidine for an outgrowth of 2-3 doublings before DNA extraction. The doublings were measured by cell count on dry runs prior to the experiment. The outgrowth at 3h starvation serves to amplify signal from live cells relative to any residual DNA from dead cells. The outgrowth at 0h also serves as a control for this process. The “Start” aliquot allowed us to identify insertion mutants with significant growth defects from the initial library generation.

#### Footprinting and Sequencing

Genomic DNA was isolated from the samples collected during the selection using the DNEasy Blood and Tissue Kit (Qiagen) and purified using Zymo genomic DNA clean & concentrate kit (Zymo Research). The isolated and purified DNA was digested with MspI and HinP1 in separate digests that are subsequently pooled. After purification again using a Zymo clean and concentrate kit, a Y-linker [91] was annealed and ligated to the DNA and the samples were purified using Agencourt AMPure beads (Beckman Coulter). The Y-linker was annealed by combining the primers with annealing buffer in a PCR tube at 90**°**C for 2 min, followed by cooling to 30**°**C at a rate of ∼2 degrees per minute. After snap cooling, the Y-linker was promptly used or frozen for future use. A second Illumina bar code was added by PCR (the first was added to the transposon fragment itself before library generation). After another bead cleanup was a second PCR amplification, which added dual indexes for sequencing. After a final bead cleanup, samples were pooled and sequenced using a NextSeq 500 sequencer (Illumina).

<u>Y-linker primers:</u>

ACTACGCACGCGACGAGACGTAGCGTC

5’phos — CGGACGCTACGTCCGTGTTGTCGGTCCTG

<u>Primers to add the second Illumina adaptor:</u>

ACACCTAACCGCTAGCACGTAATACGACTC GTGACTGGAGTTCAGACGTGTGCTCTTCCGATCTACTACGCACGCGACGAGACG

#### Survival profiling analysis

The bcl2fastq package from Illumina was used to separate the sequencing reads based on sample number and to obtain FASTQ files for each sample. Preprocessing & alignment steps clipped the Illumina adaptors & transposon sequence to identify the target sequence [92]. Quality score filtering dropped reads if the target was less than 15bps and trimmed low quality ends of reads [93]. Bowtie aligned reads to a given genome database and the total reads aligned per sample were quantified. A python script enumerated the number of footprints present within each gene, and generated data tables with the set of counts. Reads were normalized to fragments per million (FPMs). The R package rateratio.test was used to calculate significance for the differences in insertion frequencies for each gene at the 3h vs. 0h time points, aggregated at the gene level. A survival score for each gene was defined as the log_2_ fold change (LFC) of FPM at 3h/ FPM at 0h and p-values calculated assessing the significance of the observed count differences via the exact rate ratio test [94].

Two-sided exact tests and matching confidence intervals for discrete data. R Journal, 2(1), 53- 58.], and subsequently corrected for multiple hypothesis testing using the Benjamini-Hochberg method. Candidate lists were generated by collecting all genes with a significant (FDR<0.05) survival score over a threshold log_2_ fold change magnitude (+/- 0.5). For each of these genes, if its survival score calculated for FPM at 3h / FPM at “Start” had an opposite sign compared to the survival score of FPM at 3h/ FPM at 0h, they were discarded from the candidate list.

### Laboratory Evolution

Four replicates of each starting strain (*thyA^-^* in MG1655 and MDS42) were selected for survival and growth in media supplemented with 0.2μg/mL thymidine as opposed to the typically required level of 20μg/mL. The selections began by making a 100,000-fold dilution from saturated cultures of the starting strains grown overnight in media supplemented with 50μg/mL thymidine. The overnight cultures, grown to saturation before the initial inoculation, were at 37°C but once the selections began, the cells were kept aerated at 30°C to slow the rapid death process and boost the number of survivors at each transfer.

The selection process lasted for ∼50 daily transfers. All but the last 7 involved a 2-fold dilution in which the 2mL of 24h old cells and media were vortexed, 1ml was removed, pelleted, and reconstituted in 2ml fresh media. For the last 7 transfers, the dilution was increased because the number of survivors had increased; these transfers involved a 200-fold dilution of 24h cultures directly into 2mL fresh media (i.e. no pelleting).

Throughout the selection process, weekly or biweekly, the evolved batch cultures were plated onto LB/thymidine plates both at 0h and 24h post transfer in order to keep track of any large changes in survival numbers. In addition, weekly or biweekly, 15% of the 24h batch cultures were mixed with LB and glycerol and frozen at −80**°**C.

After the 50 transfers, the four heterogeneous evolved populations in each background were characterized for enhanced survival and the best surviving were spread onto LB/thymidine plates in order to isolate single colonies. These isolates were re-verified by PCR with primers flanking the *thyA* gene and, in the case of the strains in the MDS42 background, using primers flanking genes deleted in this background. Six colonies from each evolved batch culture, once verified, were grown overnight in LB+50μg/mL thymidine and frozen in LB and glycerol at −80**°**C. Isolates were characterized for increased survival in thymidine-free media. The two top surviving isolates from each background (assessed via half-life in thymidine-free media) are discussed in the main text.

### Whole Genome Sequencing

DNA was isolated using the DNeasy Blood and Tissue kit (Qiagen), then barcoded and prepared for sequencing using the Nextera XT DNA Library Prep Kit (Illumina). All samples were pooled and sequenced using a NextSeq 500 sequencer (Illumina). After the sequencing reads were demultiplexed and preprocessed, as described above for the survival profiling sequencing data, the mutations present in the evolved strains compared to the parental background were identified using breseq (version 0.23) [95].

### RNA Sequencing

The transcriptome profiling experiment mapped and quantified RNA extracted from three experimental conditions in the evolved and parental strains in the MDS42 background. Both strains were diluted 1:200 from overnight cultures into high thymidine and grown for 2h at 30°C before strains were washed twice and placed in new media. Three samples were taken for each strain: a high thymidine sample 2h into growth, a sample 30 minutes into thymidine starvation, and a control was washed twice similar to the starved sample but placed back into high thymidine and collected after 30 minutes. When the evolved strain in high thymidine was compared to the parental in high thymidine, the samples before washes and pelleting were used. When the evolved in the starved condition was compared to the evolved unstarved, the high thymidine sample post washes and pelleting was used.

After incubation with RNAprotect Bacteria Reagent (Qiagen), samples were stored at −80°C. Total RNA was made from the pellets using the Total RNA Purification Plus Kit (Norgen) according to the manufacturer’s recommended protocol for Gram-negative bacteria. Ribosomal RNA was removed from these samples using the Ribo-Zero rRNA Removal Kit (Illumina). Indexed libraries were prepared from the resulting RNA using the NEBNext Ultra Directional RNA Library Prep Kit for Illumina (New England Biolabs). Samples were pooled and sequenced on a NextSeq 500 sequencer (Illumina).

The sequencing reads were demultiplexed, and preprocessed using cutadapt and trimmomatic, as described above for the survival profiling data, and then Rockhopper (version 2.03) [32, 33] was used in two modes. First Rockhopper was used on each RNA read by itself to get quantitation and statistics on expression levels, and then it was used in paired mode on specified pairs of conditions in order to obtain significance statistics for changes in expression levels. In paired mode Rockhopper uses a negative binomial distribution to estimate the uncertainties in the read counts. The *p*-values are FDR-corrected using the Benjamini- Hochberg procedure [34]. The counts for each gene from the Rockhopper output were expressed in the normalized form, RPKM. Transcript counts were also normalized to transcripts per million (TPMs) in order to allow for more direct comparison with the survival profiling data.

### Genes With Consistent Effects Across Experimental Approaches

Gene disruptions showing a significant survival score above 0.5 and a log_2_ fold change of RNA TPMs below −0.5 in any of the three comparisons (the evolved vs. parental in starved; evolved vs. parental in unstarved; and evolved starved vs. evolved unstarved) were compiled into three lists of genes that putatively exacerbate survival during thymidine starvation. Gene disruptions showing a significant survival score below −0.5 and a LFC of TPMs above 0.5 in any of the three comparisons detailed above were compiled into 3 lists of genes that putatively alleviate survival during thymidine starvation.

#### Flow Cytometry

Fluorescence intensity was determined by fluorescence-based flow cytometry, using a BD LSR-FORTESSA instrument at the Columbia University Herbert Irving Comprehensive Cancer Center Flow Cytometry Core. Bacterial cells were grown for 2h in high thymidine at 37°C after 1:400 dilution from overnight cultures. After this pregrowth, cells were washed twice and placed into thymidine-free media (unless otherwise stated), with shaking at 37°C. Cells were stained as detailed below for the various dyes. No-dye cells were included to control for autofluorescence. Fluorescence was calculated at 1.5h and 3h into thymidine starvation. A total of ∼50,000 events for each time-point sample were determined. Gating was used to exclude cellular debris. Data were extracted using FCS Express 6.

After data acquisition, the autofluorescence was subtracted by using the data from the no-dye cells (pooled across all genotypes/conditions in a particular analysis batch) to fit a LOESS model predicting the fluorescent signal of interest as a function of the forward scatter (FSC) and side scatter (SSC) signals. The LOESS-predicted autofluorescence was then subtracted from the signal observed for each cell from the experimental samples. After the LOESS nodye-subtraction, the fluorescence signals were fit to a mixed effects model (using the lme4 R package) of the form:

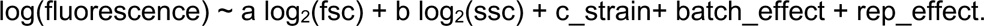

Here a and b are fixed effects indicating how the signal scales, on average, with FSC and SSC (thus explicitly accounting for changes in cell size/shape using the FSC/SSC as proxies, by factoring out any systematic changes in fluorescence that can be explained by changes in these parameters), and c_strain is a strain/condition-specific average fluorescence after accounting for the FSC/SSC effect. Batch_effect and rep_effect are random effects representing the biological replicate and day when the sample data were taken, and the inclusion of these terms allows us to factor out any systematic effects arising from day to day variations in assay behavior. The c_strain parameter was our key parameter of biological interest, and we calculated profile-based 95% confidence intervals for these parameters. In addition, the significance of differences in this parameter between biological conditions of interest was assessed using the linearHypothesis function of the R car package, against a null hypothesis that two groups had the same value of the c parameter. Only cells with both log_2_(fsc) and log_2_(ssc) greater than 8 were included in the analysis, as we found on visual inspection that events with lower values were likely simply noise.

In addition, for plotting purposes, we obtained corrected cell-level fluorescence data by calculating, for each cell, the quantity

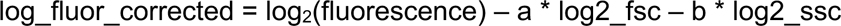

Where log2_fsc and log2_ssc refer to the specific log_2_(fsc) and log_2_(ssc) values measured for that cell. Taken together, the model yields a corrected set of fluorescence data after accounting for cell size/shape in a consistent way, as well as estimates (with confidence intervals) of the average fluorescence for each strain after correcting for size effects.

### **BCECF-AM** (Cat no. B1150, Thermo Fisher)

After pregrowth, washes, and thymidine- starvation, as detailed above, cells were incubated with 20μM BCECF in thymidine-free media (unless otherwise stated) for the last 30 minutes of the designated time period followed directly by visualization (i.e. for a 3h starvation before visualization, dye was added at 2.5h). Two channels were selected for acquiring paired sets of images: a pH-sensitive wavelength using a 488 nm laser (band pass filter 530/30) and pH-insensitive wavelength using 405 nm laser (band pass filter 525/50). In the main text and supplement, the pH-sensitive MFIs are shown.

The number of independent experiments for visualization at 1.5h: *thyA^+^*: n=6; *thyA^-^* unstarved: n=3; *thyA^-^* starved: n=5; *ΔatpF*: n=5; *ΔrecO*: n=5; *ΔspeB*: n=6; *ΔackA*: n=4; *ΔcydB*: n=5. The number of independent experiments for visualization at 3h: no dye control: 9 different strains were tested at 2-3 time points and none showed autofluorescence; *thyA^+^*: n=6; *thyA^-^* unstarved: n=8; *thyA^-^* starved: n=7; evolved MG1655: n=4; *thyA^-^*MDS42 starved: n=4; evolved MDS42: n=4; *ΔatpF*: n=6; *ΔrecO*: n=6; *ΔspeB*: n=6; *ΔackA*: n=5; *ΔcydB*: n=7; *ΔrecOΔackA*: n=3; *ΔspeBΔackA*: n=3; *ΔatpFΔackA*: n=3.

### Flow cytometry for arginine supplementation assay using BCECF

In the cases of L-arginine supplementation (and its controls) cells were incubated with 40mM L- arginine for the first 2.5h of thymidine starvation, washed twice to remove the amino acid, and incubated with dye for 30 minutes in thymidine-free media before visualization. The no-arginine control was washed and stained exactly like the cells with added arginine. The number of independent experiments for visualization at 3h: *thyA^-^* MG1655: n=5; *thyA^-^* MG1655 + Arg: n=7; *thyA^-^* MDS42: n=5; *thyA^-^* MDS42 + Arg: n=3.

### Flow cytometry for gentamicin assay for WT cells stained with BCECF

After the pregrowth and washes as outlined above, *thyA^+^*cells were treated with 1μg/mL gentamicin. After 1h (30 minutes before visualization) cells were stained with BCECF dye as outlined above. At 1.5h treatment, acidification was visualized using flow cytometry. The no drug cells were prepared, stained and visualized exactly as the drug-treated cells except for the addition of gentamicin. The number of independent experiments for visualization at 1.5h: *thyA^+^:* n=6; *thyA^+^+* gent(low): n=3.

### pHrodo^TM^ Green AM pHi Indicator

(Cat no. P35373, Thermo Fisher): 10μL of the pHrodo dye was added to 100μL of the Powerload concentrate (as specified in the pHrodo Green kit). 3.3μL of this combination was added to 300μL of cells (after pregrowth, washes, and thymidine-starvation—unless otherwise stated—as detailed above) and incubated for the last 30 minutes of the designated period at 37°C followed directly by visualization. Data were gathered at the stated times points using the following detection parameters: 488 laser, with 530/30 nm band pass filter. The number of independent experiments for visualization at 1.5h: *thyA^+^*: n=4; *thyA^-^* unstarved: n=3; *thyA^-^* starved: n=8; *ΔatpF*: n=4; *ΔrecO*: n=3; *ΔspeB*: n=6; *ΔackA*: n=3; *ΔcydB*: n=3. The number of independent experiments for visualization at 3h: no dye control: n=3; *thyA^+^*: n=3; *thyA^-^* unstarved: n=6; *thyA^-^* starved: n=12; evolved MG1655: n=8; *thyA^-^* MDS42 starved: n=8; evolved MDS42: n=3; *ΔatpF*: n=8; *ΔrecO*: n=8; *ΔspeB*: n=6; *ΔackA*: n=7; *ΔcydB*: n=7.

### Flow cytometry for arginine supplementation assay using pHrodo Green

In the case of exogenous L-arginine addition (and its controls), cells were incubated with 40mM L-arginine for the first 2.5h of thymidine starvation, washed twice to remove the L-arginine, and the cells were incubated with dye for 30 minutes in thymidine-free media before visualization. The no-arginine control was washed and stained exactly like the cells with added arginine. The number of independent experiments for visualization at 3h: *thyA^-^* MG1655: n=4; *thyA^-^* MG1655 + Arg: n=6; *thyA^-^* MDS42: n=3; *thyA^-^* MDS42 + Arg: n=3.

### Flow cytometry for Gentamicin assay for WT cells stained with pHrodo Green

After the pregrowth and washes as outlined above, *thyA^+^*cells were treated with 1μg/mL gentamicin for the lower dose (Fig 4I) or with 4μg/mL gentamicin for the higher dose (S10 Fig). 30 minutes before visualization cells were stained with pHrodo Green as outlined above. At 1.5h treatment, acidification was visualized using flow cytometry. The no drug cells were prepared, stained and visualized exactly as the drug-treated cells aside from the addition of gentamicin. The number of independent experiments for visualization at 1.5h: *thyA^+^*: n=5; *thyA^+^+* gent(low): n=3; *thyA^+^*+ gent(high): n=2.

### Peroxy Orange 1, Tocris (Cat no, 49-441-0, Fisher Scientific)

After pregrowth and washes as detailed above, cells were placed into thymidine-free media along with 5μM dye and incubated, shaking, at 37°C, for the designated amount of time before visualization. Data were gathered at the stated times points using the following detection parameters: 561 nm laser, 585/40 nm band-pass filter. The number of independent experiments for visualization at 1.5h: *thyA^+^*: n=3; *thyA^-^* unstarved: n=4; *thyA^-^* starved: n=3; *ΔatpF*: n=3; *ΔrecO*: n=3; *ΔspeB*: n=3; *ΔackA*: n=3; *ΔcydB*: n=3. The number of independent experiments for visualization at 3h: no dye control: n=3; *thyA^+^*: n=6; *thyA^-^* unstarved: n=6; *thyA^-^* starved: n=10; evolved MG1655: n=3; *thyA^-^*MDS42 starved: n=10; evolved MDS42: n=3; *ΔatpF*: n=16; *ΔrecO*: n=5; *ΔspeB*: n=12; *ΔackA*: n=14; *ΔcydB*: n=9; *ΔrecOΔackA*: n=5.

### Flow cytometry for exogenous H202 control

at 1h thymidine starvation, 1mM hydrogen peroxide was added to cells stained with Peroxy Orange dye as detailed above and incubated for 30 minutes at 37°C before visualization. The number of independent experiments for visualization at 1.5h: *thyA^-^* unstarved: n=4; *thyA^-^* unstarved + H_2_0_2_: n=5; *thyA^-^*: n=3; *thyA^-^*+ H_2_0_2_: n=3.

#### Flow cytometry for the gentamicin assay for WT cells stained with Peroxy Orange

After the pregrowth and washes as outlined above, *thyA^+^* cells were stained with Peroxy Orange dye as outlined above. Treated cells were also supplemented with 1μg/mL gentamicin for the lower dose (Fig 4K) or 4μg/mL gentamicin for the higher dose (S10 Fig). At 1.5h treatment, ROS accumulation was visualized using flow cytometry. The no drug cells were prepared, stained and visualized exactly as the drug-treated cells except for the addition of gentamicin. The number of independent experiments for visualization at 1.5h: *thyA^+^*: n=5; *thyA^+^*+ gent(low): n=3; t*hyA*^+^+ gent(high): n=2.

#### Calculating MICs for gentamicin

The MIC for *thyA^-^* (parental) and *ΔackA* were calculated using the Liofilchem® MIC Test Strip for Gentamicin (cat. Number 22-777-677; 0.016-256 μg/mL) on LB plates supplemented with thymidine.

## Acknowledgements

We wish to thank past and present members of the Tavazoie laboratory for critical discussion and comments on the manuscript. We are also grateful to Siu-Hong Ho and Wei Wang at the Columbia University Herbert Irving Comprehensive Cancer Center Flow Cytometry Core for help using the Fortessa instrument. Research reported in this publication was supported by the National Institutes of Health under award numbers R01AI077562 (to S.T.), R35GM128637 (to P.F.), and training grant 1F31GM108419- 01 (to A.K.)

## Supporting Information

**S1 Fig.**
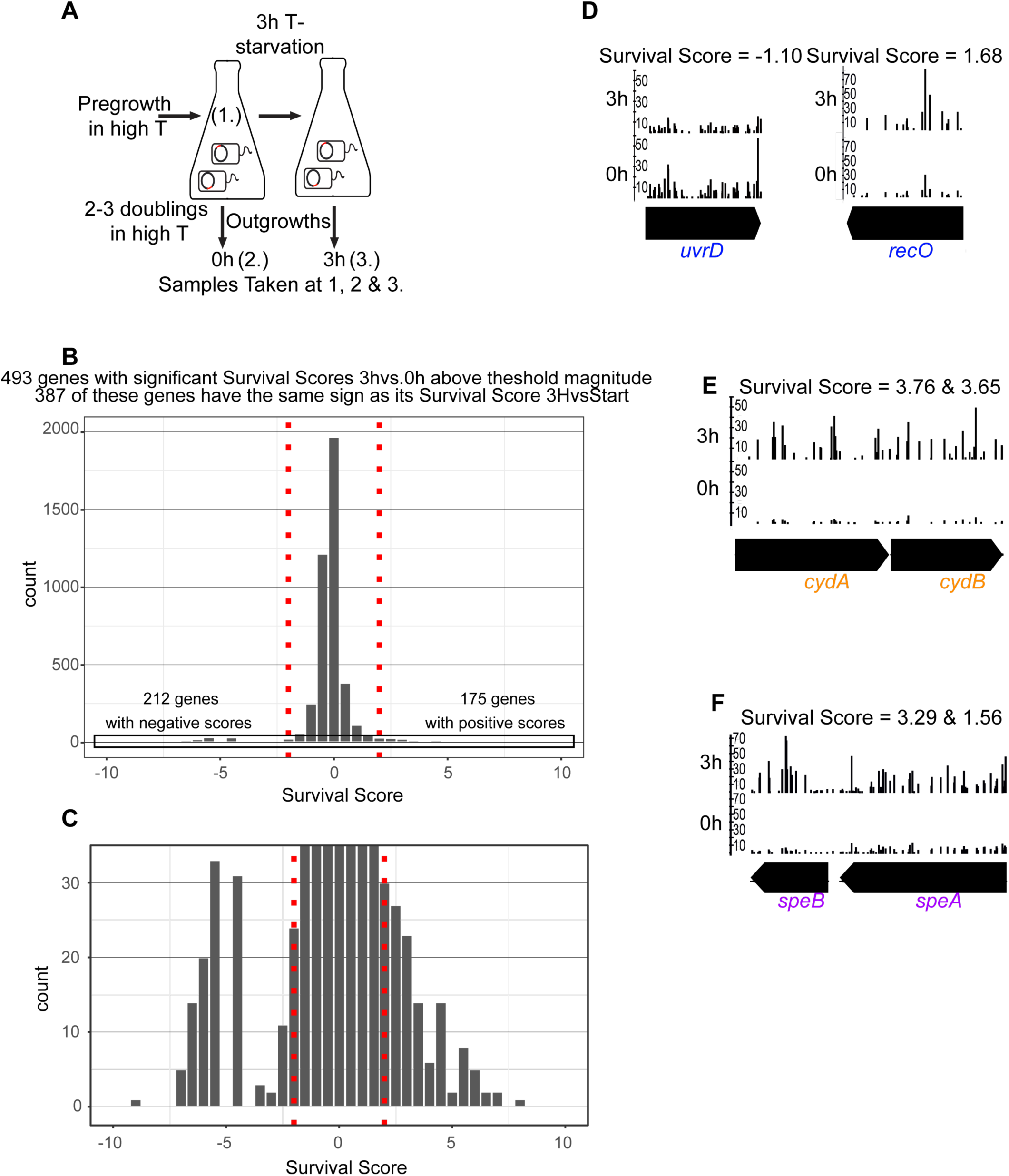
Survival profiling uncovers known and novel contributors to TLD. (A) Survival profiling time points, with and without outgrowths. At 0h starvation and 3h starvation, an aliquot of the transposon library was placed in high thymidine for 2-3 doublings in order to amplify signal from living cells over residual signal from dead cells. An aliquot was also taken at the start of the selection, before the 0h outgrowth. (B) The distribution of survival scores and generation of candidate lists. 493 genes were above a threshold log_2_ fold change magnitude (shown by red dashed line) and had a *q*-value<0.05. Significance was calculated on the ratio of FPM at 3h vs. 0h using the rate ratio test and was corrected using the Benjamini & Yekutieli method. Any genes on this list that had an opposite sign from the survival score for the 3h vs. “Start” were discarded, yielding a total of 387 candidate genes. (C) is an inset for the area pictured in the rectangle in panel (B). (D-F) Transposon insertions visualized using the Integrated Genome Browser. The frequency of each insertion site is shown in read counts per million. (D) Frequency of transposon insertions along the length of two previously known TLD contributors at 0h and after 3h thymidine starvation. A deletion in *uvrD* is known to sensitize cells to TLD and a deletion in *recO* is known to alleviate TLD. (E) Enhancements of transposon insertions within the *cydAB* operon. (F) Enhancements of transposon insertions within the *speAB* operon.

**S2 Fig.**
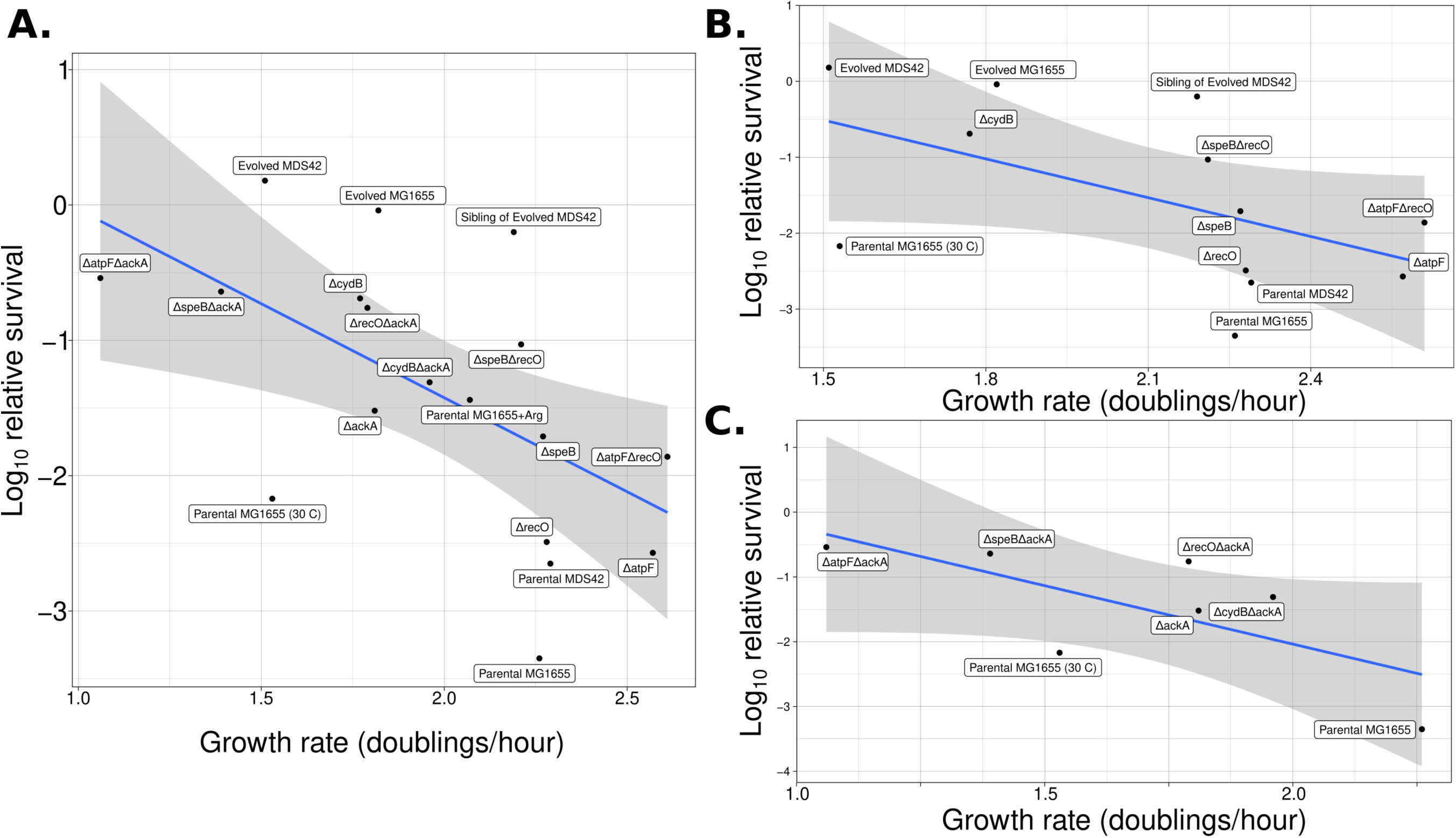
The correlation of growth rate in high thymidine rich defined media vs. relative survival at 3h in zero thymidine media. A. All strains. B. All strains minus the *ackA* strains. C. *ackA* strains only. See Methods under Growth Rate and Death Assays.

**S3 Fig.**
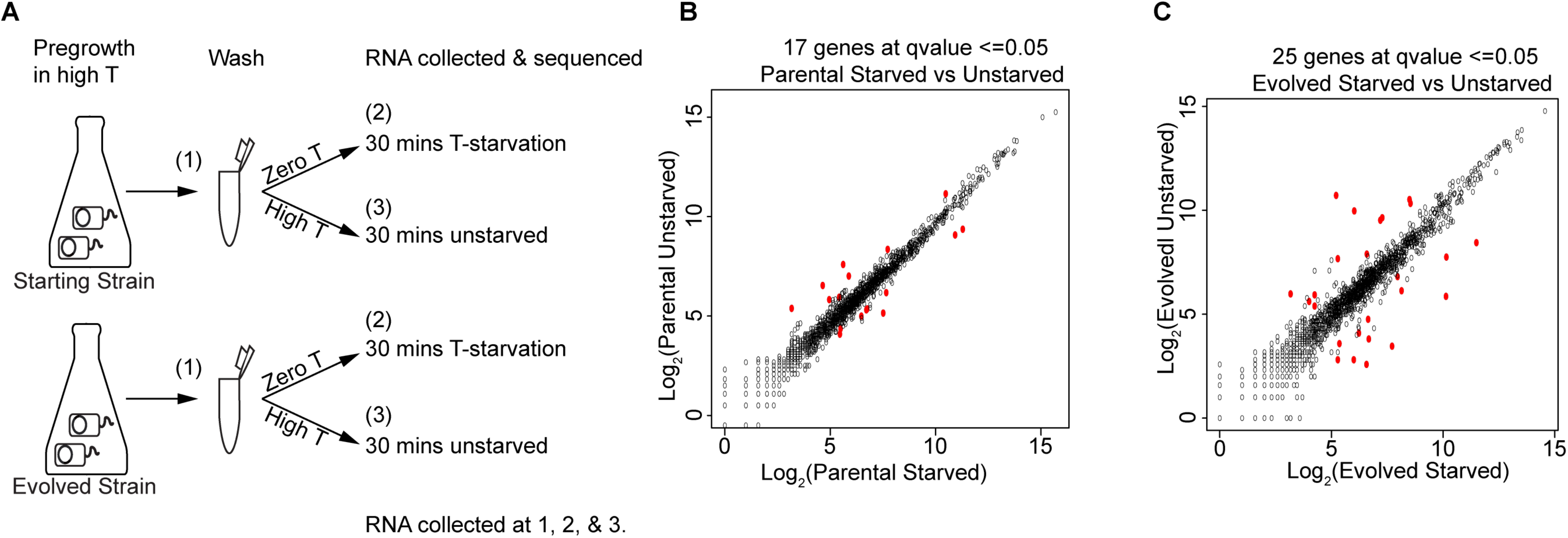
Transcriptional responses of TLD-sensitive and TLD-resistant strains. (A) Experimental setup of RNA sequencing of parental and evolved strains in the MDS42 background. All steps were performed at 30°C (the temperature at which laboratory evolution was conducted). Pregrowth was for 2h, and RNA was collected both before pelleting (1) and 30 minutes after pelleting and transfer into thymidine-free media (2). RNA was also collected 30 minutes after pelleting and placement back in high thymidine as a control (3). (B-C) Log2 expression of genes in two comparisons showing significant differentially expressed genes in red. The 17 and 25 significant differentially expressed genes can be found in S4D&C Figs, respectively. (B) Parental starved vs. unstarved. (C) Evolved starved vs. unstarved.

**S4 Fig.**
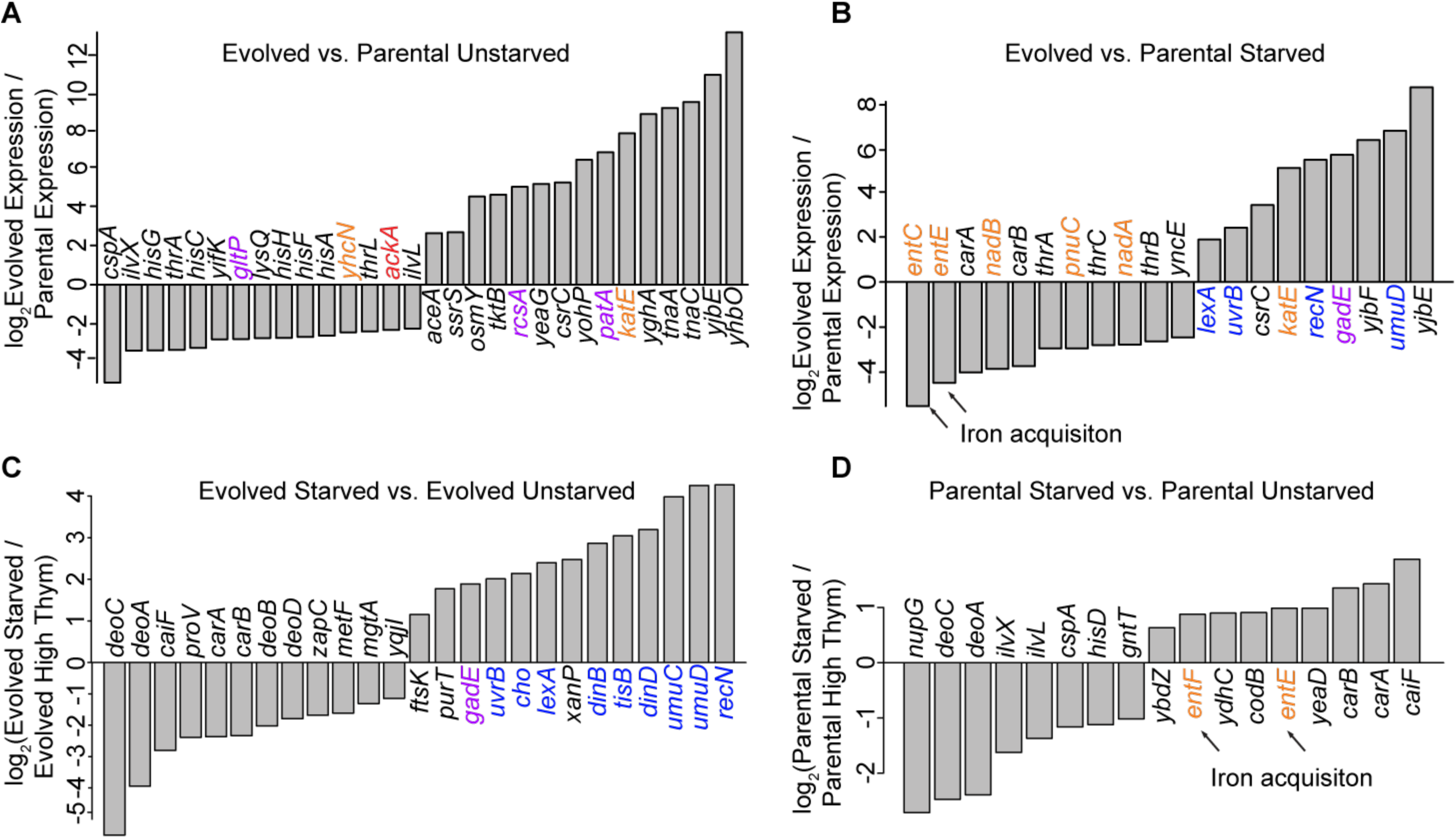
Gene-level analysis of transcriptional responses. (A-D) Significant differentially expressed genes in various comparisons. See S6 Table for the *q-*values. The LFCs of RPKMs are visualized here. (A) Significant differentially expressed genes in the evolved vs. the parental in the unstarved condition. (B) Significant differentially expressed genes in the evolved vs. parental 30 minutes into thymidine starvation. (C) Significant differentially expressed genes in the evolved starved vs. evolved unstarved. (D) Significant differentially expressed genes in the parental starved vs. parental unstarved. As in the main text, purple represents genes involved in putrescine / glutamate / arginine metabolism or their associated amino acid decarboxylation acid resistance systems; orange represents genes involved in ROS; red represents genes involved in acetate dissimilation; and blue represents genes involved in DNA replication/repair.

**S5 Fig.**
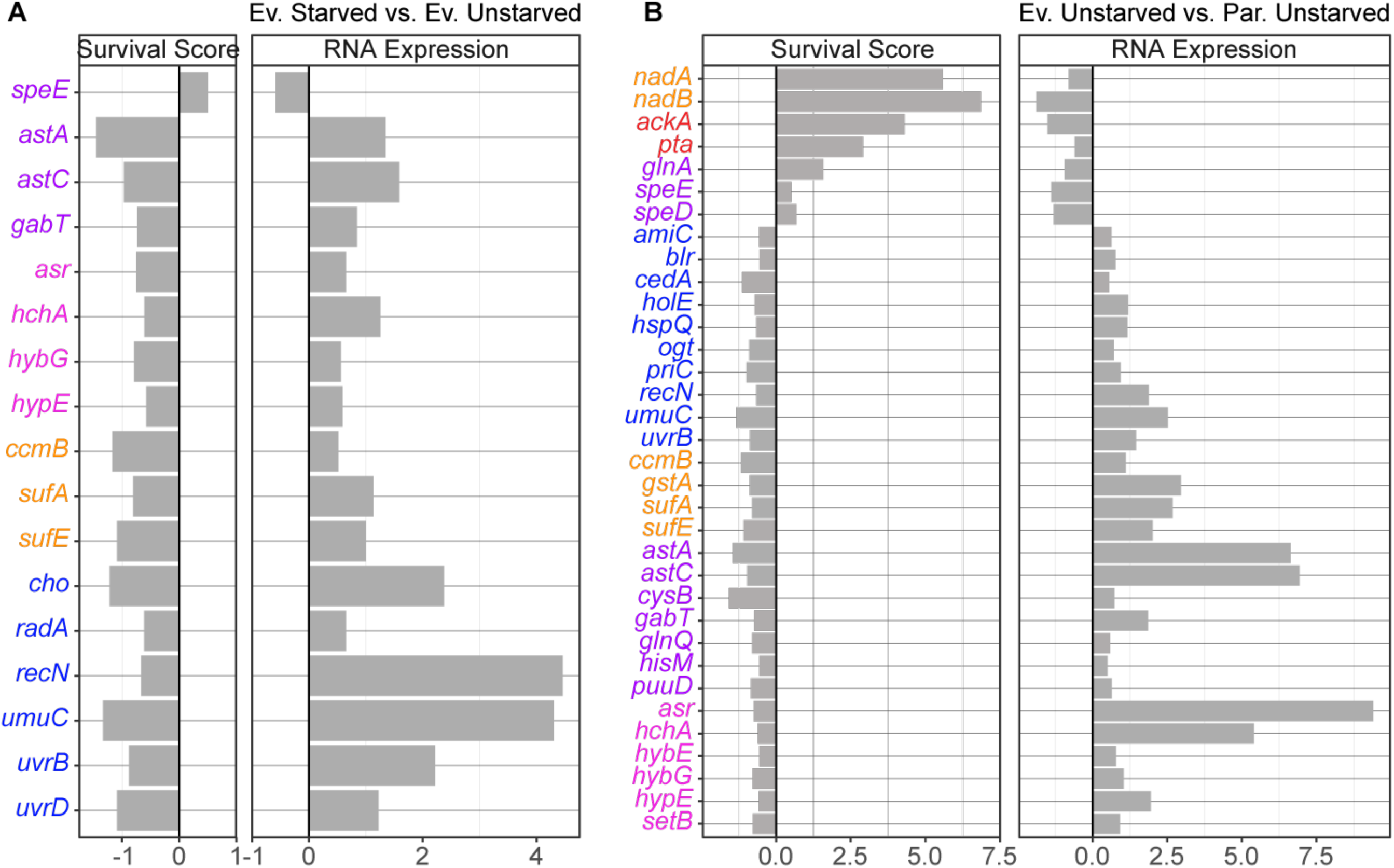
Genes in DNA replication/repair, Electron transport chain and/or ROS accumulation, and pH homeostasis pathways with consistent genetic and transcriptional effects across experimental approaches. (A-B) The left-hand column shows survival scores from the survival profiling experiment and the right-hand column shows LFC of mRNA expression (TPM) from the transcriptome profiling experiment. (A) Survival scores with the LFC of RNA expression in evolved starved vs. evolved unstarved. The remaining genes showing consistent effects can be found in S8 Table. (B) Survival scores with the LFC of RNA expression in evolved unstarved vs. parental unstarved. The remaining genes showing consistent effects can be found in S9 Table. Colors are used to represent recurring pathways as described in S5 Fig. Pink represents genes involved in proton translocation (or sequestration) systems or in the acid stress response.

**S6 Fig.**
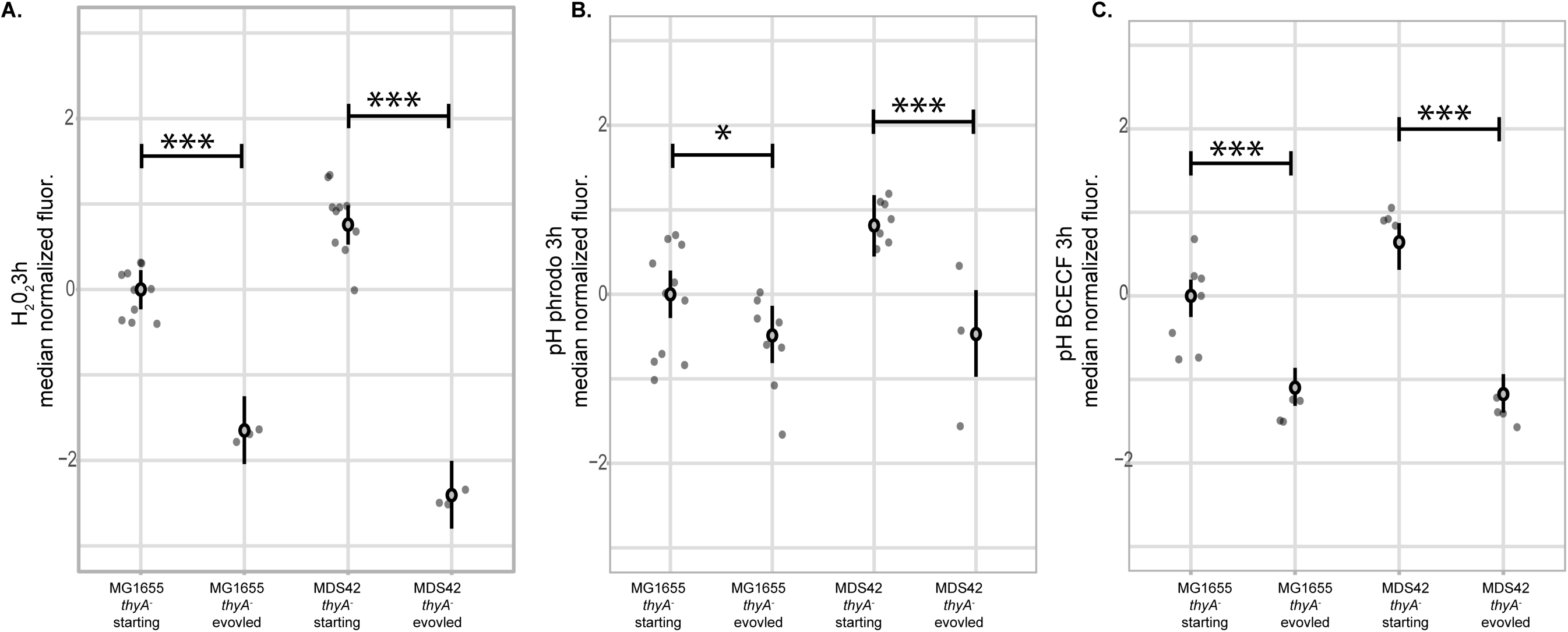
Parental strains in two genetic backgrounds show evidence of ROS accumulation and acidification at 3h thymidine starvation while the evolved TLD-resistant strains do not. (A-C) Adjusted fluorescence measured using flow cytometry. All strains are in thymidine-free media. (A) Adjusted fluorescence measured at 3h for strains dyed for hydrogen peroxide using Peroxy Orange. (B) Adjusted fluorescence measured at 3h for strains dyed for pHi using PHrodo Green. (C) Adjusted fluorescence measured at 3h for strains dyed for pHi using BCECF-AM. See Fig. 3 caption for definitions of plotted intervals and significance tests.

**S7 Fig.**
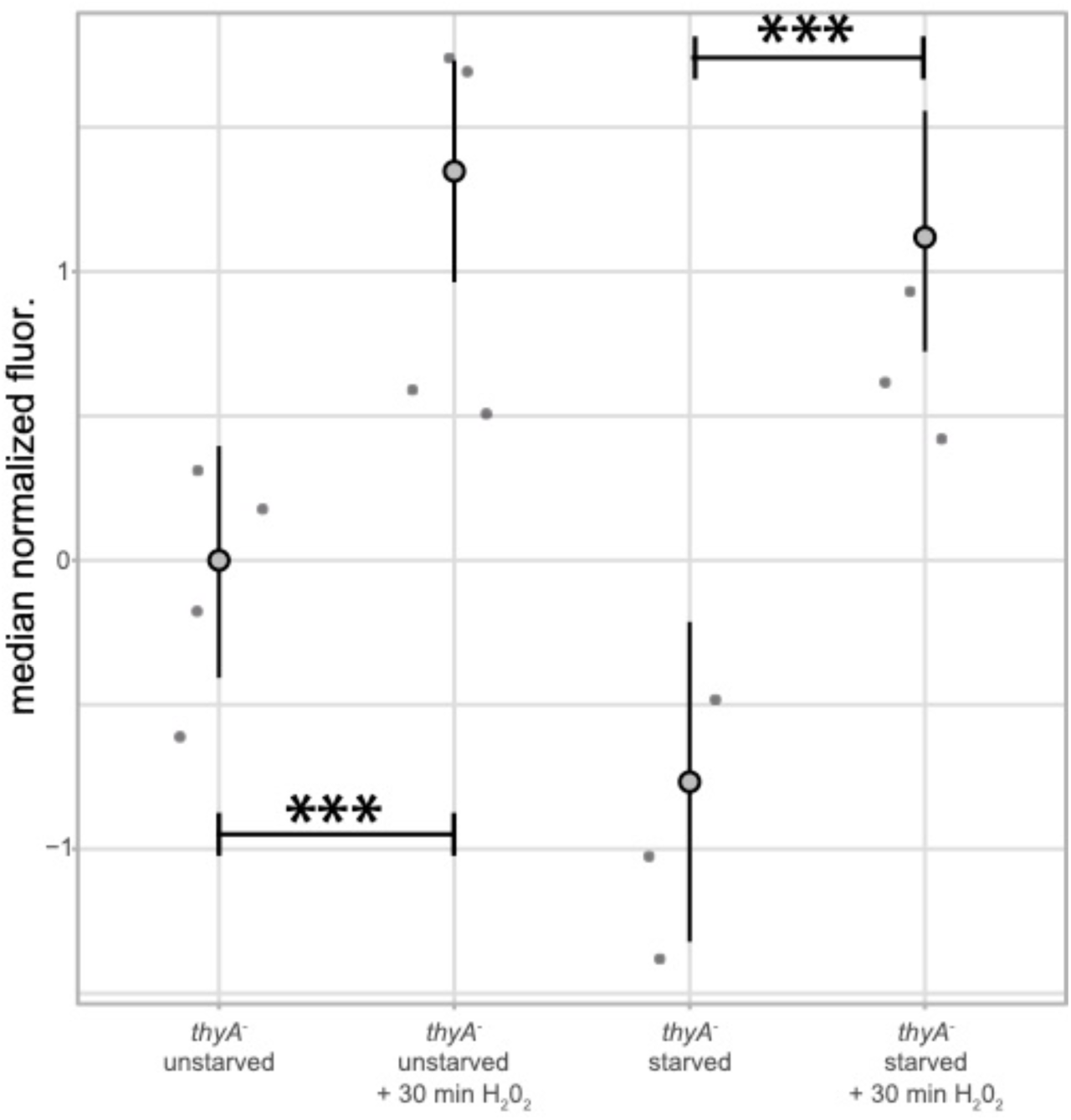
*thyA^-^* strain shows evidence of ROS accumulation at 1.5h only when exogenous H_2_0_2_ is added. Adjusted fluorescence measured using flow cytometry of *thyA^-^* cells stained with Peroxy Orange with and without a 30-minute incubation with 1mM H_2_0_2_. The two plots on the left show adjusted fluorescence at 1.5h in high thymidine media and the two plots on the right show adjusted fluorescence at 1.5h thymidine starvation. See Fig. 3 caption for definitions of plotted intervals and significance tests.

**S8 Fig.**
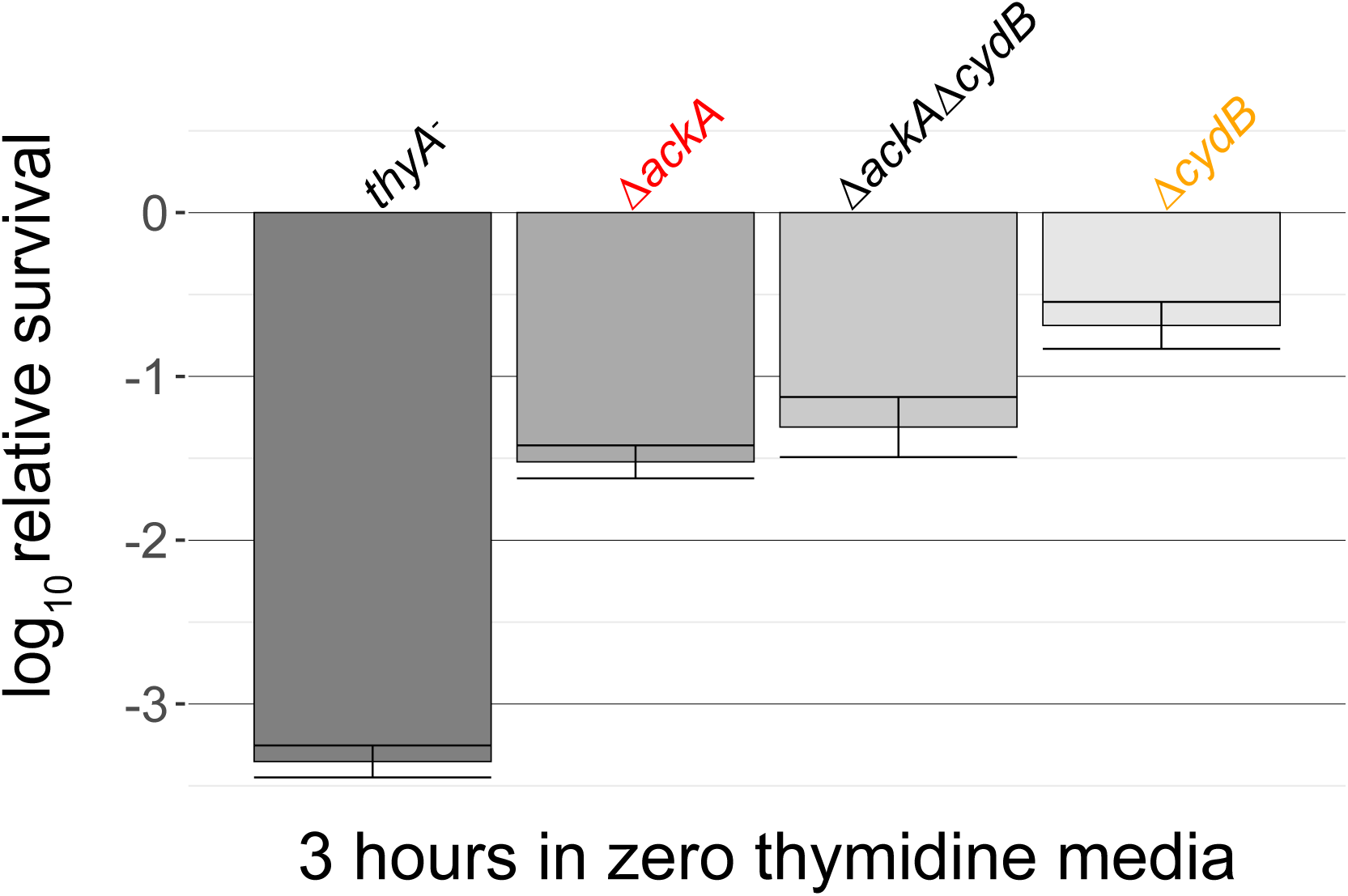
The *ΔackAΔcydB* double mutant does not show increased survival. Survival of the parental strain and knockouts in the MG1655 background at 3h thymidine starvation. Relative survival was measured for at least three independent experiments, with error bars representing standard error of the mean.

**S9 Fig.**
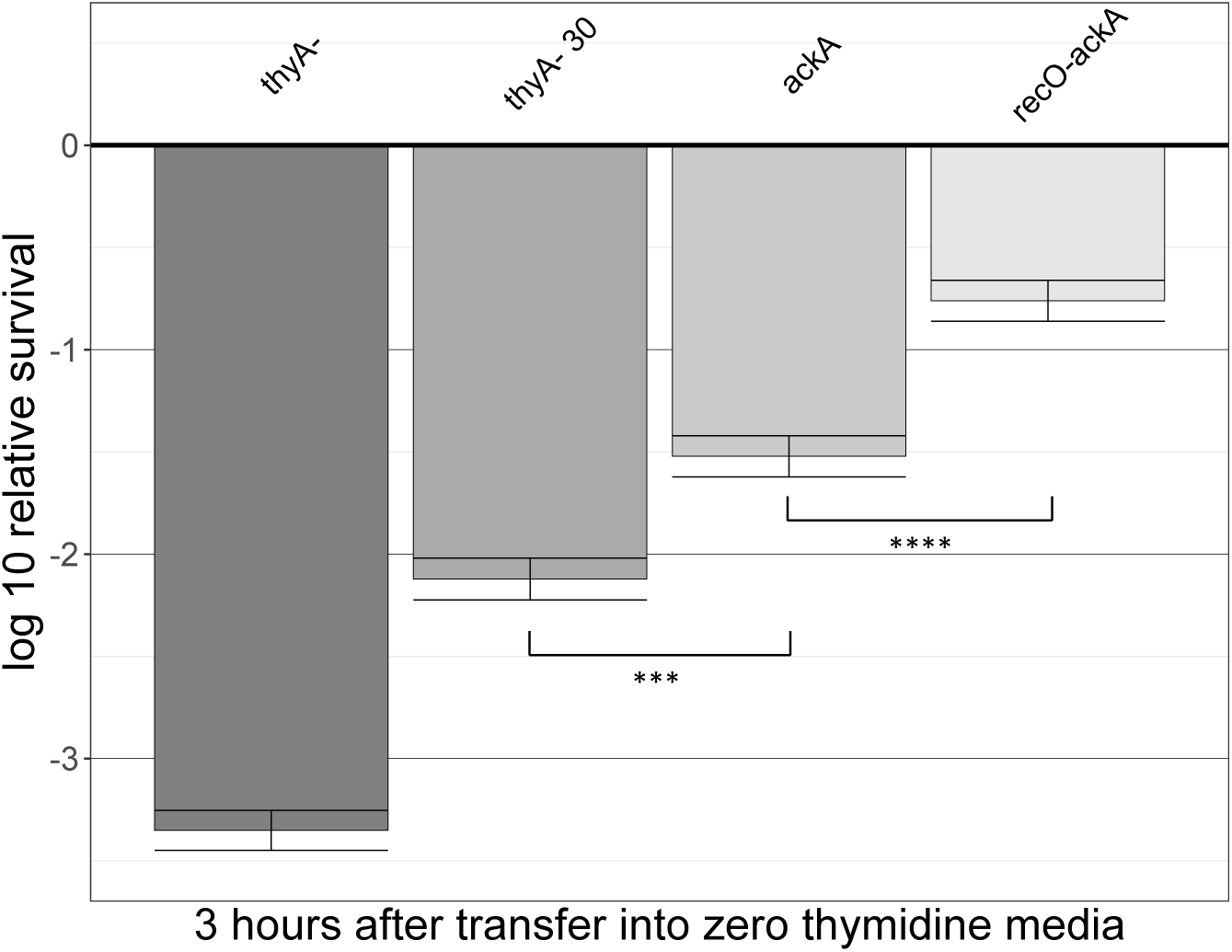
Growth at 30 degrees has less effect alleviating TLD than the *ackA* and the *recO+ackA* mutants. Survival of the parental strain (*thyA^-^*) and knockouts in the MG1655 background at 3h thymidine starvation at 37° vs 30°. Relative survival was measured for at least three independent experiments, with error bars representing standard error of the mean. T-test significance for *ackA* vs. *ackA recO* = 8.24 x 10^-05^ and for the *thyA^-^* at 30° vs. ackA = 5.29 x 10^-4^.

**S10 Fig.**
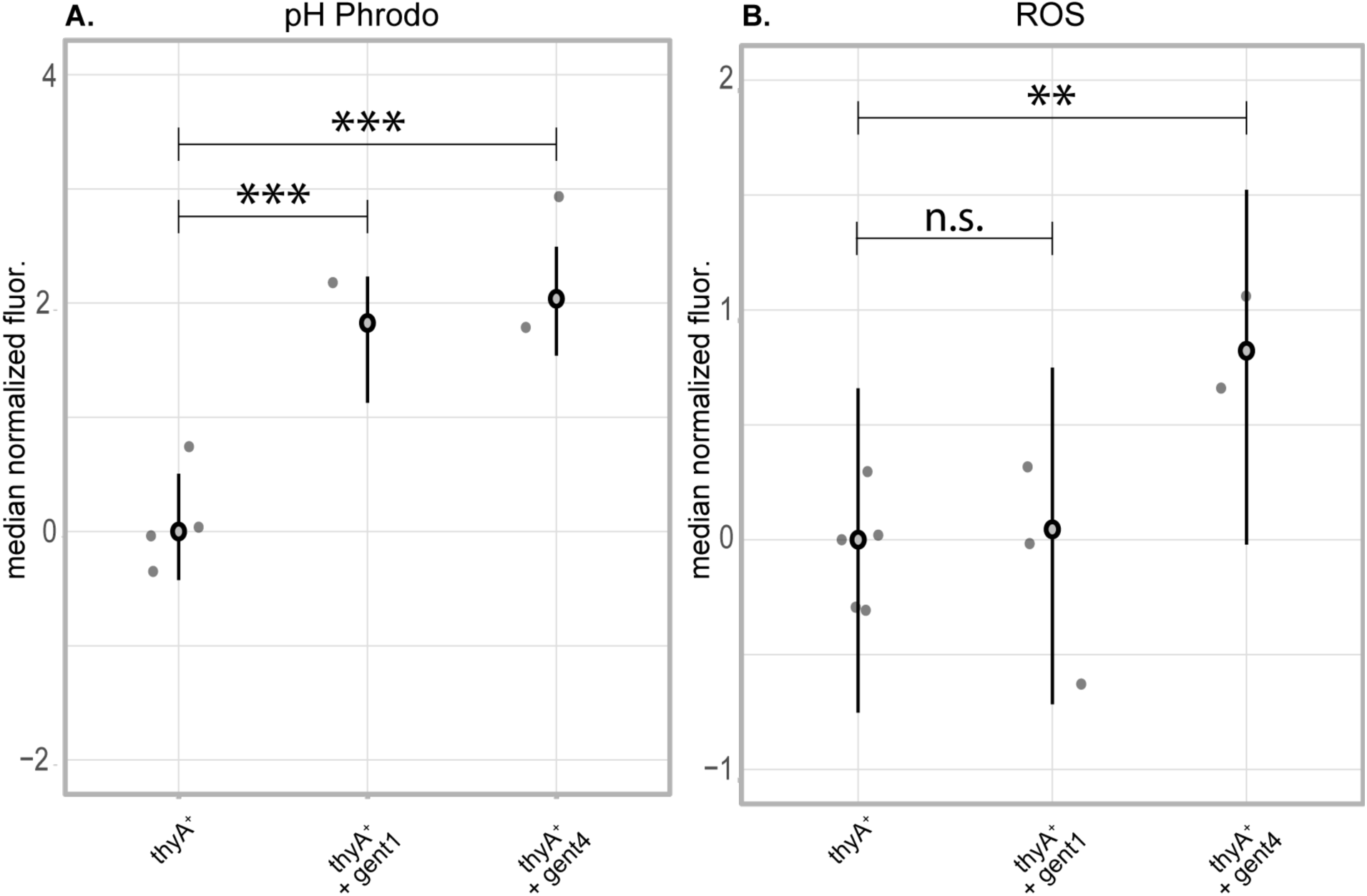
At higher concentrations of gentamicin, both acidification and ROS accumulation are observed. Plots of adjusted fluorescence of wild type (*thyA^+^*) cells measured by flow cytometry at 2h with and without 1μg/mL or 4μg/mL gentamicin using pHrodo Green for pHi (A) or Peroxy Orange for ROS (B). See Fig. 3 caption for definitions of plotted intervals and significance tests.

## Supplementary Tables

S1 Table: Survival scores and significance for the survival profiling of a saturated transposon library generated in the MG1655 thyA^-^ genetic background.

S2 Table: Survival profiling candidate genes in which disruptions exacerbate killing. S3 Table: Survival profiling candidate genes in which disruptions alleviate killing.

S4 Table: Mutations in the TLD-resistant evolved isolate in the MG1655 background.

S5 Table: TPMs of RNA expression of parental and evolved strains in the MDS42 genetic background.

S6 Table: RPKM, TPMs and q-values of significant differentially expressed genes from the transcriptome profiling experiment.

S7 Table: Candidate genes that show consistent effects in genetic and transcriptional responses in evolved starved vs. parental starved.

S8 Table: Candidate genes that show consistent effects in genetic and transcriptional responses in evolved starved vs. evolved unstarved.

S9 Table: Candidate genes that show consistent effects in genetic and transcriptional responses in evolved unstarved vs. parental unstarved.

